# Humans are colonized by many uncharacterized and highly divergent microbes

**DOI:** 10.1101/113746

**Authors:** Mark Kowarsky, Joan Camunas, Michael Kertesz, Vlaminck Iwijn De, Winston Koh, Wenying Pan, Lance Martin, Norma Neff, Jennifer Okamoto, Ron Wong, Sandhya Kharbanda, Yasser El-Sayed, Yair Blumenfeld, David K. Stevenson, Gary Shaw, Nathan D. Wolfe, Stephen R. Quake

## Abstract

Blood circulates throughout the entire body and contains molecules drawn from virtually every tissue, including the microbes and viruses which colonize the body. Through massive shotgun sequencing of circulating cell-free DNA from the blood, we identified hundreds of new bacteria and viruses which represent previously unidentified members of the human microbiome. Analysing cumulative sequence data from 1,351 blood samples collected from 188 patients enabled us to assemble 7,190 contiguous regions (contigs) larger than 1 kbp, of which 3,761 are novel with little or no sequence homology in any existing databases. The vast majority of these novel contigs possess coding sequences, and we have validated their existence both by finding their presence in independent experiments and by performing direct PCR amplification. When their nearest neighbors are located in the tree of life, many of the organisms represent entirely novel taxa, showing that microbial diversity within the human body is substantially broader than previously appreciated.

---

In many high-throughput sequencing experiments there exists a set of reads that are unable to be aligned to any existing reference databases. These reads do not have any known sequence homology, and after sequencing artifacts are properly accounted for they are typically interpreted as a signature of novel organisms^1^. Applying this approach to environmental metagenomics has led to the discovery of many new phyla, expanding knowledge of the diversity of the tree of life^2^, and large human microbiome studies such as the Human Microbiome Project^3^ and MetaHIT^4^ have characterized many previously unknown taxa at easily accessible body sites. However, those projects targeted specific niches such as the gut or skin, and therefore do not detect organisms residing in other body sites or those possessing low abundances. Here, we take advantage of the fact that blood is a medium that samples virtually the entire body and collects molecules – including DNA – released by the organisms which colonize humans in all body sites.

The existence of circulating nucleic acids in blood has been known since the mid-20th century^5^, but only in the last few years has the advent of high-throughput sequencing led to clinical diagnostics based on these nucleic acids (also known as cell-free DNA or RNA), including detecting fetal abnormalities^6^, transplanted organ rejection events^7^ and signatures of cancers^8^. It is not only human cells that shed their nucleic acids into the blood; other life forms such as viruses, bacteria, fungi also release their DNA and RNA into the blood, providing a source that can help determine the presence of infectious disease^9^ or measure alterations of the virome due to pharmacological immunosuppression^10^. There is roughly an order of magnitude more non-human cells than nucleated human cells in the body^11,12.11,12^ Combining this with the average genome sizes of a human, bacterium and virus (Gb, Mb, kb respectively) suggests that approximately one percent of DNA by mass in a human is derived from non-host origins. There is ample evidence that blood is able to sample this set of foreign DNA molecules and we define this non-host DNA as the ‘infectome’. Previous studies by us and others have shown that indeed about 1% of cell free DNA sequences appear to be of non-human origin, but only a small fraction of these map to existing databases of microbial and viral genomes. This suggests that there is a vast diversity of as yet uncharacterized microbial diversity within the human microbiome.

We analysed the infectome of 1,351 samples from 188 patients in four longitudinally sampled cohorts - heart transplant (HT) - 610 samples [76 patients]; lung transplant (LT) - 460 samples [59 patients]; bone marrow transplant (BMT) – 161 samples [21 patients]; pregnancy (PR) - 120 samples [32 patients]) - and discovered that the majority of assembled sequences are derived from previously unidentified organisms. For example, we found numerous novel anelloviruses in immunocompromised patients, which represent a doubling of identified members in that viral family. Over two thirds of the sequences are bacterial and the majority are most similar to proteobacteria; however many large contigs can only be classified at the phylum or superkingdom level. We also found numerous novel phages throughout the population. Multiple independent analyses confirm the existence of these novel sequences.

## Results and discussion

We sequenced a total of 37 billion molecules from the 1,531 samples of cell-free DNA, of which 95% of reads passed quality control. Of these an average of 0.45% did not align to the reference human genome (GRCh38) (figure 1A left), in line with our expectations of the DNA sources in the body. Only about a percent of these reads could be identified in a curated infectome database of almost 8,000 species lying within 1,651 genera of known bacteria, viruses, fungi and some eukaryotic pathogens (figure 1A, third from left). This miniscule fraction of reads encompasses the *known* infectome. Less than 1,800 known species (800 known genera) are observed across all samples. The rarefaction curve of species prevalence quickly plateaus and the species abundance distribution has only a slight positive skew (figure S1). These both indicate that the number of known species we measure has saturated^13^ and deeper or broader sequencing of cell-free DNA from humans is unlikely to substantially increase the richness of known species.

**Figures 1.**
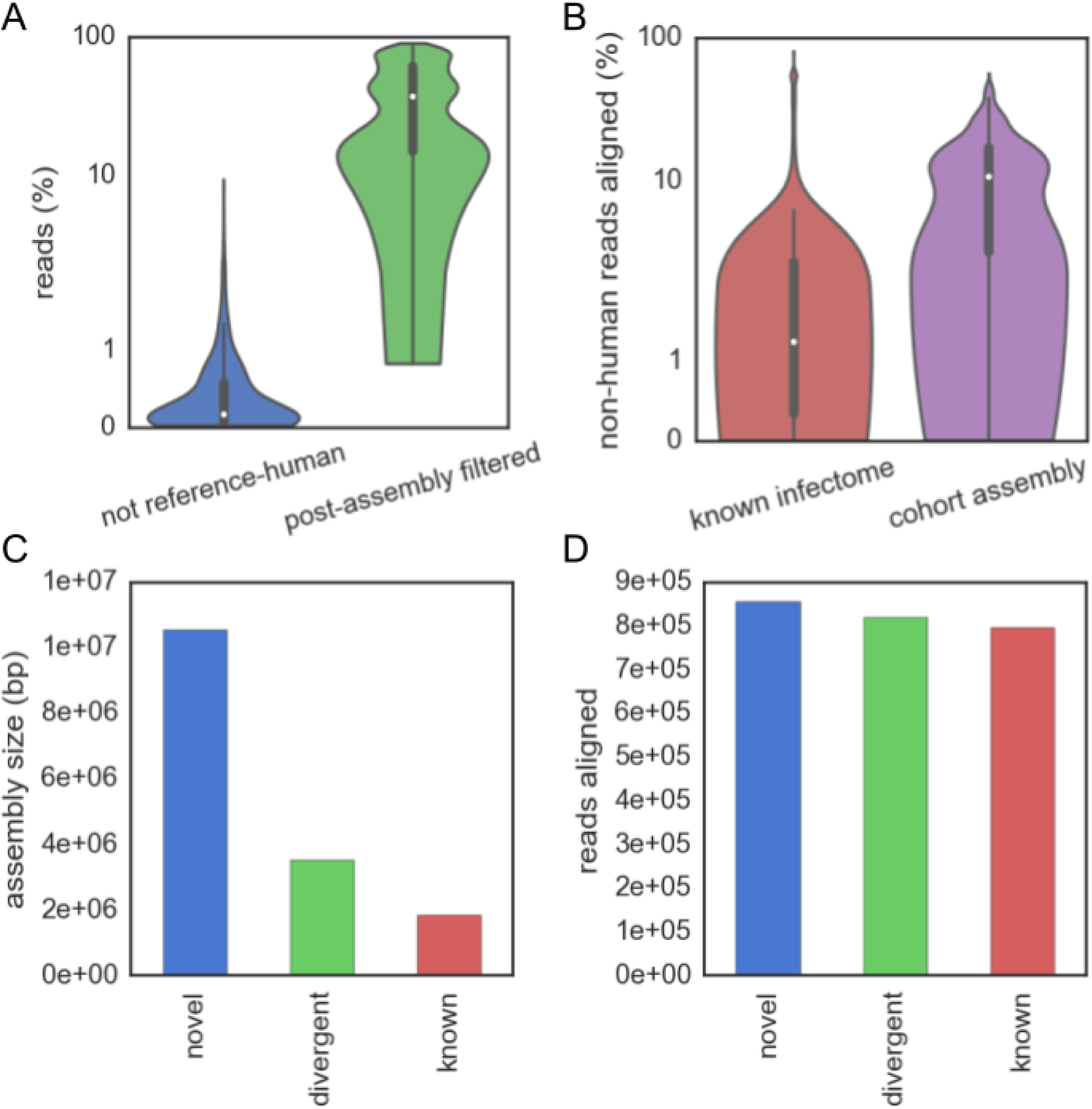
A-B) Violin plots showing the distributions of the percentage of reads associated with various stages of the assembly pipeline. From left: A) reads that did not align to the reference human sequence; reads that were not removed during post-assembly cleanups of additional human-like or low-complexity sequences; B) non-human reads that aligned to a curated infectome database; non-human reads that aligned to the cohort assemblies. The white dot is the median value and thicker black bar the interquartile range. C) The total size of novel, divergent and known contigs in the cohort assemblies. D) Number of reads aligning to novel, divergent and known contigs in the cohort assemblies.

We performed *de novo* assembly on the remaining non-human reads to uncover new species in the dark matter of cell-free DNA. The construction of assemblies used an iterative approach. Non-human reads were assembled on a per-sample basis and reads that aligned to low-complexity or human-derived contigs were removed. This process of assembly and cleaning was repeated for remaining reads, pooled first by patient and then by cohort, and resulted in a total assembly of 40 Mbp. Over 25 megabases of low complexity or human derived contigs were removed (15% of reads) (figure 1A, second from left), many of which were identified as human microsatellites or BAC/FOSMID clones (supplementary file *removed_sequence_table.tsv*). The iterative assembly process also functions to capture more reads in each stage as the number of reads pooled together increases (figure S2). The cohort assemblies constructed 7,190 contigs larger than 1 kbp and 131 larger than 10 kbp. Compared to the proportion of reads that map to known organisms, an order of magnitude greater fraction of reads map to the cohort assemblies (figure 1A, right half).

To select for contigs likely to originate from uncharacterized genomes, a series of filtering steps was applied. The first two filters enrich for contigs that have a high gene content (i.e. predicted genes span at least 60% of bases)) and low homology at both the nucleotide and protein level to any previously known sequence (figures S3 and S4, i.e. BLAST alignments span less than 20% of bases and an average gene identity of less than 60%). Application of these filters results in the selection of 4,353 contigs over 1 kbp. Two additional steps removed contigs that have homologies in expanded microbiome sequence databases. First, we aligned all assemblies from phases II and III of the HMP and blacklisted the 47 contigs that had matches. As many of the HMP assemblies are of previously known organisms or had been deposited in the NCBI nt database, the BLAST coverage filter encompasses most of the HMP filtered contigs (figure S5). The second filter used the web service Onecodex^14^ to remove contigs with homologies in their database.

To control for potential contaminants from the extraction columns, we prepared six sequencing libraries using the same protocol used in the plasma-based cell-free DNA samples, but instead of plasma utilized either water or DNA extracted from a human cell-line. Filtered reads were aligned to the assembled contigs, resulting in blacklisting an extra 114 contigs due to their suspected presence in control samples. An additional step to check for contaminants was performed using the non-host reads from 300 cfDNA samples obtained from nonhuman primate plasma. No contigs were observed at a level above 1 read per kilobase in more than 158 samples, with 75% of contigs observed at this level in less than 27 samples. The highly variable and non-ubiquitous expression of these contigs in primate samples indicates that these are not common contaminants from the lab or kits.

After all the filters, a total of 4,187 contigs remain, which can be further reduced to 3,761 novel candidates after merging contigs with significant overlaps. For later comparisons, a further 773 contigs are classified as ‘known’ (>80% BLAST coverage and >1 kbp) and a further 598 as divergent (>1 kbp and neither known nor novel). The majority of assembled bases are novel, with 11 Mbp assembled compared with 3.5 Mbp of known or divergent contigs (figure 1B). The number of reads aligning is similar for all classes of contig (figure 1C). The lower coverage of novel contigs may have kept them hidden from previous analyses, and only by pooling many samples together were these sequences found^15^.

Among the 3,761 novel contigs we found 14 with ribosomal proteins, none of which resembled the 16S ribosomal subunit. This coupled with the fact that many of the sequences are viral-like led us to conclude that a taxonomic classification based on 16S rRNA sequences is unfeasible. Thus, the taxonomic position was estimated based on choosing the lowest common ancestor of the majority of genes with homology, based on the gene’s taxonomic assignment from alignment. For known contigs, this method agrees with the best nucleotide alignment of a contig at the species or lower level over 85% of the time, indicating the suitability of this approach for taxonomic classification. Contigs with both bacterial and phage genes had their taxonomic placement refined by ignoring the phage genes if that made the assignment more specific. To visualize the taxonomic diversity, a ‘solar system’ was constructed for all novel contigs (figure 2) Individual dots are contigs with the rings representing the taxonomic levels; their radial position within the ring is proportional to the average gene identity (inner is high, outer is low); and the angle their particular superkingdom or phylum. The 844 yellow contigs placed on the outermost ring cannot confidently be placed in any superkingdom based on gene homology because for almost all of them the predicted genes have no known homology. Over two thirds of sequences appear to be bacterial, the majority of these are most similar to proteobacteria. Most longer contigs (>5kb) are bacterial or prophage-like and can only be placed at the phylum or superkingdom levels.

**Figure 2.**
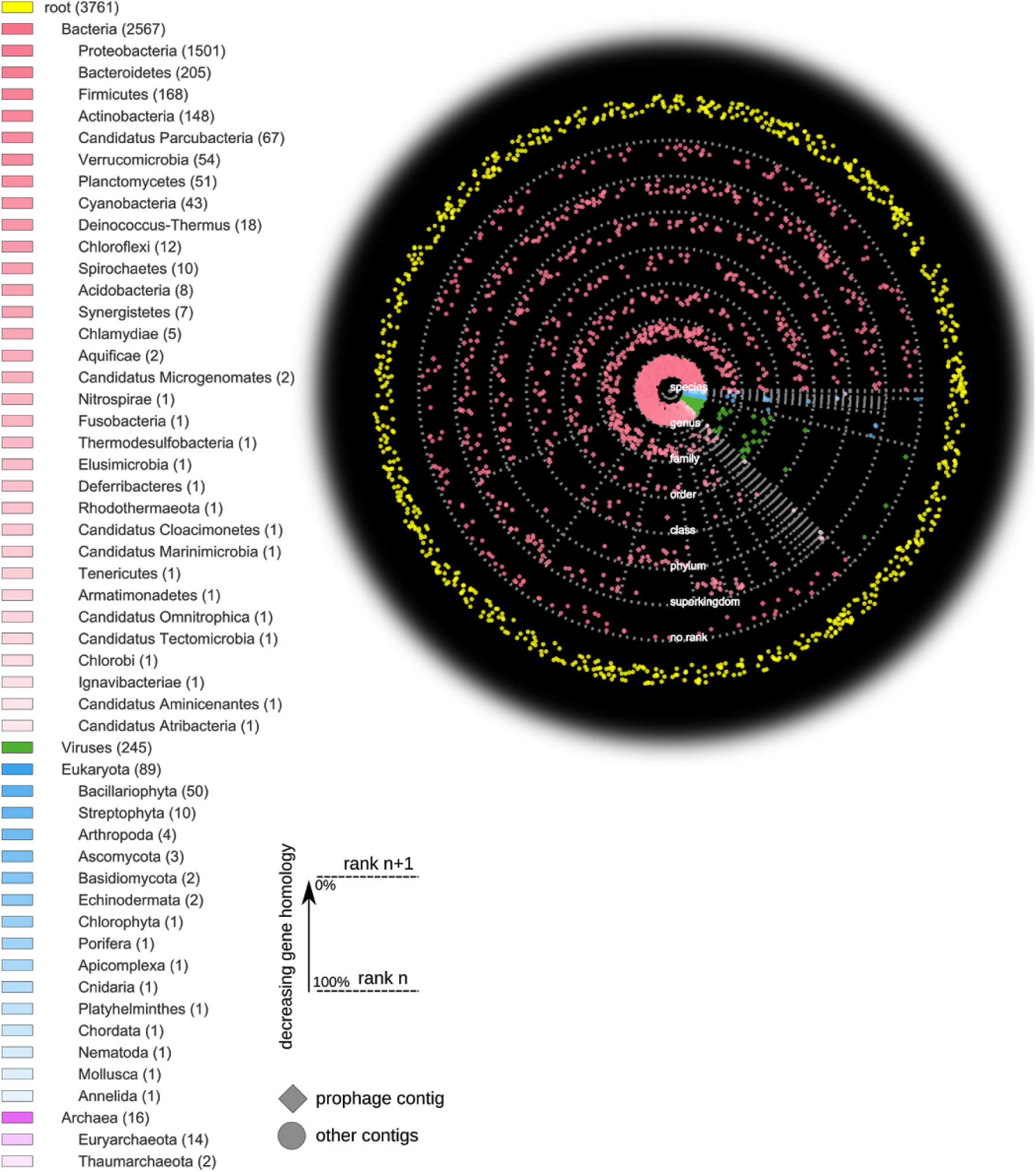
Solar system plot of all novel contigs (>1 kbp) and a taxonomic tree showing the assigned levels (root, superkingdom, phylum). Each ring represents a different taxonomic level, with the intra-ring radius representing the average gene homology. Points are randomly assigned within sectors based on their superkingdom or phylum, as represented by the grey dotted spokes. The yellow contigs outside the last ring could not be assigned to any superkingdom, the radial scattering is to help illustrate the density of contigs. Colours correspond to superkingdom, intensity to the the phylum. Potential prophages have diamond shapes.

The novel sequences indicate that the infectome’s species richness is much higher than is estimated from known species. If we count all the taxa the contigs are assigned to, the number of taxa of known and divergent contigs is ~20% of the total number of species observed in the curated infectome database. Conservatively counting contigs assigned to higher levels only once (e.g., only count ‘proteobacteria’ once), we observe over 1,000 novel taxa. If the discovery rate is similar to known and divergent contigs, this gives a range of 1-5 thousand new species (i.e. a 50-250% increase over known species).

The distributions of average gene identity within each rank are clearly distinct for novel, divergent and known contigs (figure S6), which is reflected in their solar system plots (figures S7 and S8). Although the divergent and known contigs are fewer in number, they are much more precisely placed, both in their taxonomic level and their average gene identity (reflected in the points being placed closer to the rank rings in the solar system). As more sequences are identified or classified, we expect many of the novel contigs in the outer orbitals to be attracted towards the center. The novel contigs are not restricted to individual samples or patients and most appear ubiquitously across the cohort populations. Each novel contig is observed in a median of 51 patients and the median number of novel contigs observed in each patient is 924 (also see histograms in figures S9 and S10). Even though elements of the human microbiome have been well-studied by deep sequencing from particular body sites, it is evident that other niches accessed by plasma contribute substantial novel diversity which we report here for the first time.

From the 2,917 placed novel contigs, 276 (9%) correspond to novel viral sequences which are predominantly either phages or torque teno viruses (TTVs). Distinguishing between a phage and its bacterial host is difficult with short sequences, as they both are prone to incorporate each other's genes. Indeed, of the 523 contigs containing phage genes, 333 also have bacterial genes. Nonetheless, identifying these is important as the contigs with the most predicted genes are all phage or prophage candidates. Half of the genes have no homology for the top 15 such contigs, with some (HT_node_2, HT_node_16, cluster_12) having over three quarters of genes without matches. Of the identified genes, their closest homologies are commonly hypothetical proteins. These two facts conspire to make functional annotation unfeasible at this time. The non-hypothetical genes present on the phage contigs include DNA primases, phage tail proteins, terminases, capsid proteins and DNA polymerases I and III. Many of their bacterial genes are from species within the proteobacteria phylum, with some from actinobacteria (HT _node _11, LT_node_6) or containing genes from a flavobacterium (bacteroides phylum) phage. Additional sequences were searched for in the recently published global virome from the Joint Genome Institute^16^. Only 21 of the unmerged contigs had matches over 50% of their lengths, and these derived from just six of JGI’s scaffolds. Three of these are from Coloradan soil samples, one each from a wastewater bioreactor, a freshwater lake and from the upper troposphere. Our contigs matching to these are mostly best assigned as a cyanobacteria phage or unclassified Siphoviridae. The phage-like contigs were highly prevalent across patients and exhibited only minor clustering by cohort (figure S11). These phages are likely to be associated with bacteria in the background microbial flora. Given that these contigs are observed ubiquitously in our data (77-287 samples), and seen in the JGI data only in a few exotic environments, we hypothesize that they are bona fide members of the human virome and their presence in the JGI data is due to coincident discovery of related environmental species.

In contrast, the novel TTVs show strong clustering amongst cohorts (figure S12) on the basis of immune system status. TTVs encompass Anellovirus families and are known to be enriched in immunocompromised patients.^10^ Their detection here represents another validation of the taxonomic assignment based on gene homology. A phylogenetic tree was built using multiple sequence alignment of these contigs along with reference sequences from previously characterized anelloviruses (figure 3). Both reference and *de novo* assembled contigs can be clustered into three distinct classes: a class of non-human infecting reference anelloviruses (green), a set of novel anelloviruses that have only 35-48% sequence similarity with any reference sequence (yellow), and the rest which we identify as new species or subspecies of existing TTVs. This work has almost doubled the total number of anelloviruses found in humans including a new potential genus of human-infecting anelloviruses. Therefore assembly based metagenomic methods can uncover vast amounts of prevalent diversity even in known viral families with numerous reference sequences.

**Figure 3.**
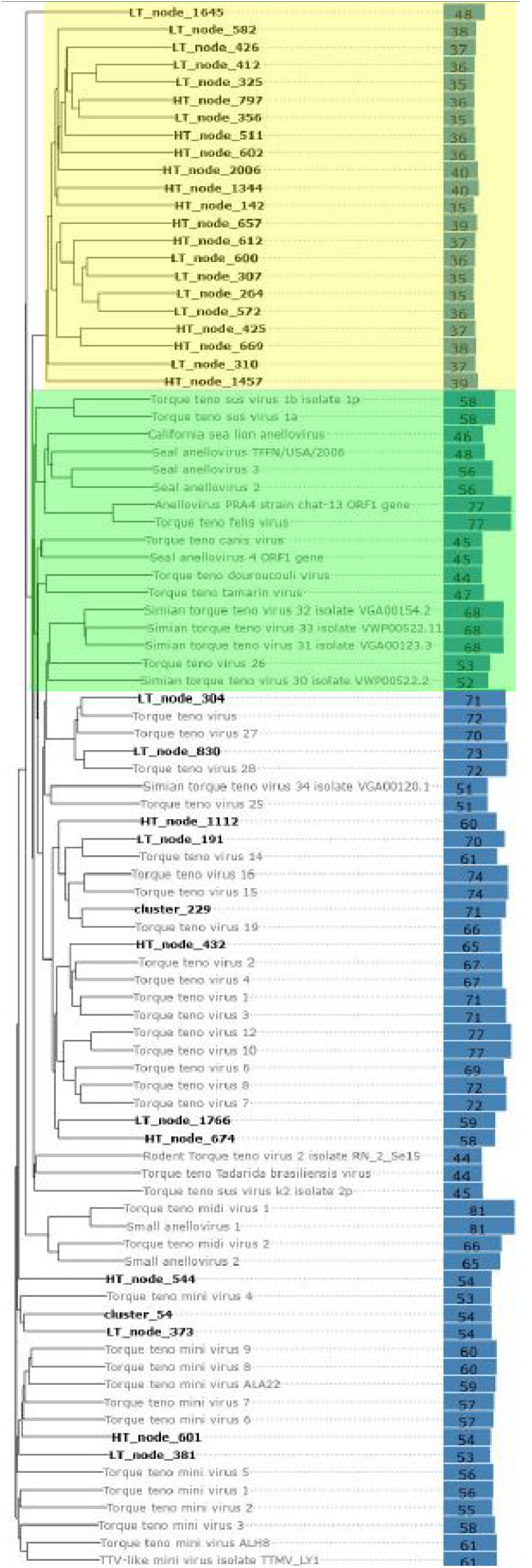
Phylogenetic tree of reference and *de novo* assembled (bold) torque teno viruses (TTVs). Yellow sequences a class of contigs divergent from known torque teno viruses (TTVs), green are known TTVs in animals. On the right are blue bars (with numbers) indicating the percentage identity of the nearest (non-self) reference sequence.

Many novel sequences cannot be classified based on their gene level homologies. There is a clear trend in the reduction of both the proportion of genes identified and the degree of similarity to known genes as we compare the known, divergent and novel contigs (figure 4, figures S13A-C). This lack of homology is the reason there are 840 unrooted contigs unable to be placed into any existing superkingdom (the yellow points in the outer solar system of figure 2). The majority of these (n=739) have four or fewer predicted genes. Only four of the unrooted contigs have any identified genes and most of those that are identified are of unknown function (full table in supplementary file *unrooted_contigs.txt*).

**Figure 4.**
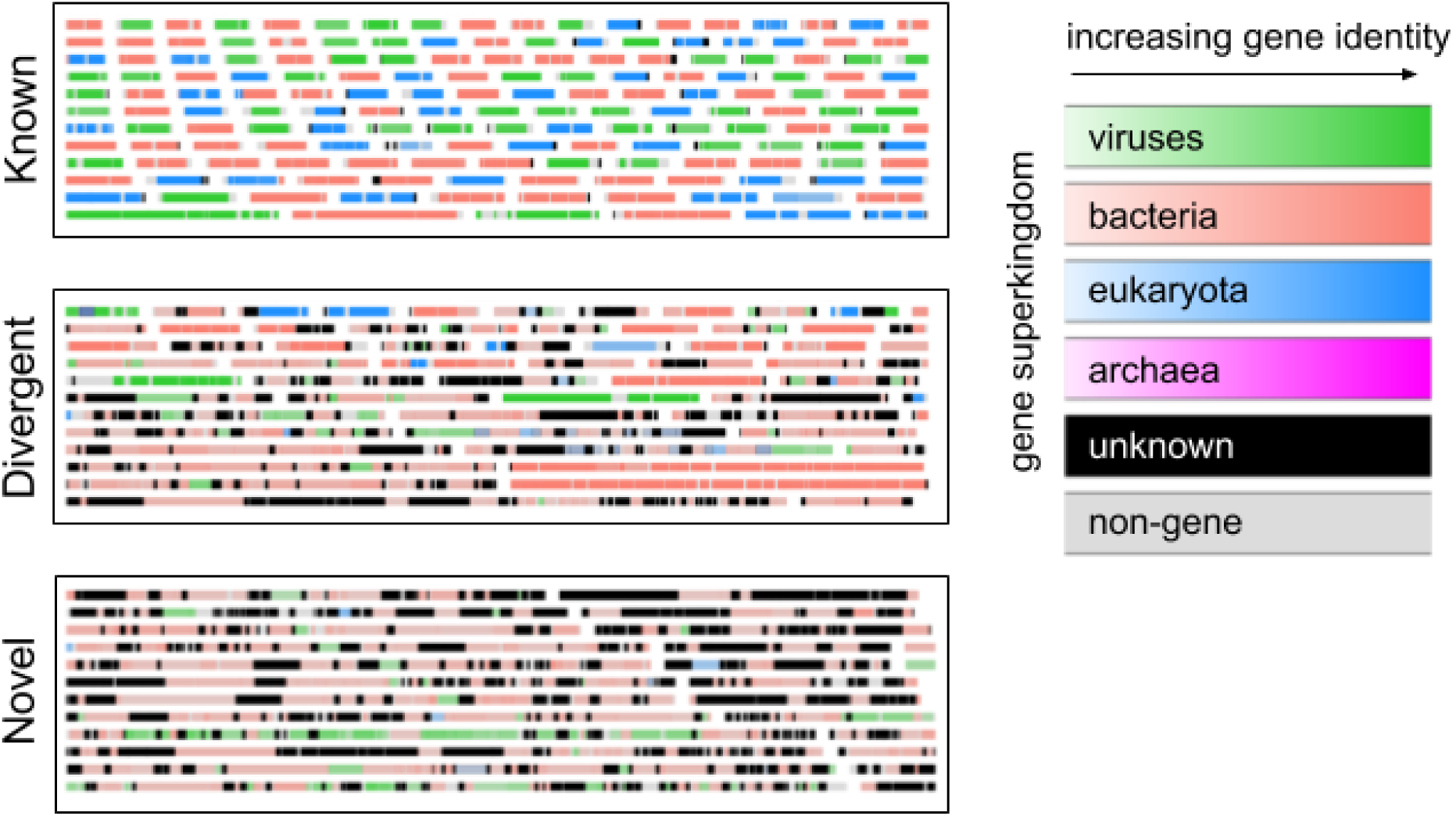
Examples of contigs and their gene assignments for known, divergent and novel contigs. Width of the box is equivalent to 60 kkbp. Genes are coloured by superkingdom, with saturation being proportional to the gene identity.

**Figure 5.**
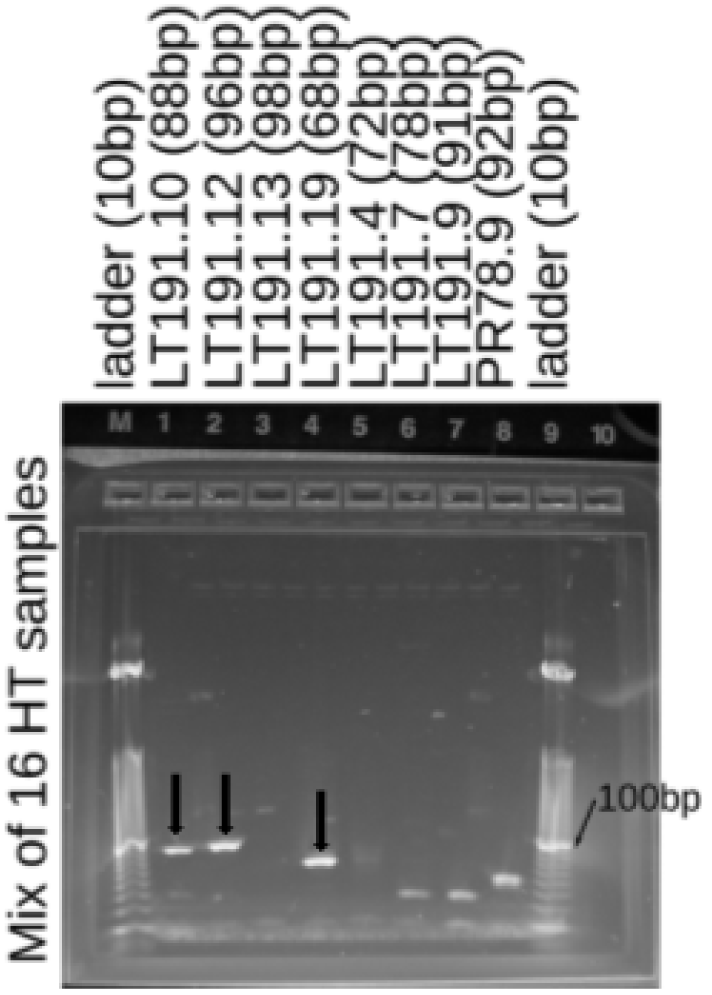
Gel showing the presence of three segments of LT_node_191 (a novel torque-teno virus) in the cfDNA deriving from the heart transplant recipients (Eight patients, two samples per patient).Demultiplexed gels and no template controls are in figure S19.

As alignment based methods failed to identify any genes from the remaining unrooted contigs, an estimated superkingdom assignment was made based on the codon usage of genes^17^. Comparing all genes with and without homology shows that these classes possess significantly different codon usages (p < 1e-3 after Bonferroni correction, Mann-Whitney U test, figure S14), which is not merely an artifact of differential GC bias (figure S15). An adaptive boosting classifier trained on known genes allowed us to bin all except 16 of the unrooted contigs into single or multiple superkingdoms (figure S16). The vast majority (n=640) are bacteria, and similarly to the rooted contigs, many others (n=87) have viral and bacterial genes and are treated as phage or prophages. There are hints that horizontal gene transfer across superkingdoms may be prevalent in biological dark matter, with 106 unrooted contigs having diverse sets of genes. Despite this substantial taxonomic distance from known species, we were able to reliably and consistently assign the novel contigs to taxonomic locations on the tree of life.

As validation that the novel contigs are not merely contaminants or assembly artifacts, we performed a series of additional tests of their existence. The fragmented nature of cell-free DNA prevents direct PCR amplification of large regions of the contigs, so instead we (i) screened external datasets and samples for novel sequences at the read level, (ii) used bioinformatic approaches to assess assembly quality and (iii) independently measured the presence of novel contig sequences by PCR in additional samples.

Plasma aliquots from a lung transplant recipient in our cohort were independently prepared and sequenced in another laboratory using an alternative library preparation protocol^18^. We downloaded the reads from the NCBI sequence read archive and processed them identically in the pipeline followed by aligning them to the database of novel contigs. The two sets of different preparations exhibit concordant rankings of the high-abundance contigs (figure S17). We also downloaded data generated by another group^19^ from completely independently collected samples and extraction methods and compared it to our pregnancy cohort (figure S18). Although these samples had only slightly more than a tenth as many non-human reads as our samples, there were still reads which mapped to the novel contigs discovered in our experiments. These external datasets provide additional evidence that the novel contigs are not local laboratory contaminants and that their presence is robust to different sample preparation protocols.

The quality of assembly was assessed using information from the iterative assembly approach. Alignments between contigs from different levels created a graph structure, allowing us to investigate the additive structure derived from pooling more samples together. Under ideal circumstances, contigs assembled at lower stages should be wholly contained at higher stages and they shouldn’t end up orphaned (i.e. having no homology to a later stage’s contig). Only 3,301 of the 25,765 sample contigs (over 500 bp, non-human, not low-complexity) are orphaned when compared with the patient contigs, Over 3,000 of these are from single-end read samples, and are ignored by the assembler at later stages as the longer paired end reads tend to be more informative. These mostly had high BLAST coverages (10,768 had coverage >20%) and encompassed the following genera most frequently: Lactobacillus (5,975), Streptococcus (1,449), Saccharomyces (1,315), Leuconostoc (1,051) and Lactococcus (610). Likewise to the sample to patient case, the vast majority of the remaining contigs (>85%) derive from patients who only have single-end sequencing. If we look in the other direction, ie at the lower contigs contained within the novel contigs, the majority of them derive from a single patient’s assembly (with some reads recruited from others), and only 387 weren’t assembled at any lower stages. The increased complexity of assembling due to more pooling does not appear to reduce the power to assemble what is present or to produce strong evidence of chimeric sequences.

An additional check of the pipeline with a synthetic control dataset was performed to test if interesting sequences are prematurely removed. This was comprised of 8,068 curated genomes from the NCBI of viruses, bacteria and fungi from which high-quality synthetic 100 bp reads at 25 bp intervals were created. Only 28 genomes (supplementary file *control_remove_pipeline.tsv)* had more than 50% of their reads removed in the pipeline steps leading up to assembly. These include various plasmids and enterobacteria phages (expected to be filtered in UniVec), simian virus 40, a few candidate bacterial species and two variants of the human papillomavirus. Less than 10% of genomes had more than 10% of their reads removed, and those that were removed are sequences known to be in the cleaning or subtraction databases, indicating that the subtraction steps are unlikely to remove interesting non-human sequences.

We performed PCR testing to verify the presence of novel contigs in 16 previously unsequenced samples derived from the plasma of eight heart transplant recipients. As the fragmented nature of cell-free DNA prevents the amplification of large fragments, we designed primers against numerous short (60-100 bp) templates (all gels in figure S19). Sequencing libraries known to express the templates targeted were used as positive controls. Eight primers for one of the novel TTVs (LT_node_191) and eight primers unrooted contigs (HT220, HT552, HT76) were used. One of the eight patients had evidence at both time points for four templates from the TTVs, three additional patients had evidence for one TTV-derived template. None of the unrooted templates were seen in these samples. This was not a surprising fact, as the median of maximum number of reads (over samples) is 21, in contrast, the TTVs have a median value of 180 reads sequenced. Only through sequencing many foreign molecules from hundreds of samples were the unrooted contigs (and other novel ones) able to be found.

## Conclusion

Deep sequencing of cell-free DNA from a large patient cohort revealed previously unknown and highly prevalent microbial and viral diversity in humans. This demonstrates the power of alternative assays for discovery, and shows that interesting discoveries may lurk in the shadows of data acquired for other purposes. Many megabases of new sequences were assembled and placed in distant sectors of the tree of life. With deeper sequencing and targeted sample collection, we expect numerous new viral and bacterial species to be discovered in the circulating nucleic acids of organisms that will complement existing efforts to characterize the life within us. Novel taxa of microbes inhabiting humans, while of interest in their own right, also have potential consequences for human health. They may prove to be the cause of acute or chronic disease that, to date have unknown etiology, and may have predictive associations that permit presymptomatic identification of disease.

## Methods

### Sequencing and preprocessing

Plasma was extracted from whole-blood samples as previously described^6^ and stored at -80C. Prior to sequencing, they were thawed and circulating DNA was extracted using the QIAmp Circulating Nucleic Acid Kit (Qiagen). Libraries were prepared using the NEBNext DNA Library Prep Master Mix Set for Illumina with standard Illumina indexed adapters (IDT) or using an automated microfluidics based platform (Mondrian ST; Ovation SP Ultralow Library Systems). The Agilent 2100 Bioanalyzer (High Sensitivity DNA Kit) was used to characterize libraries, and they were sequenced on the Illumina platform (HiSeq 2000, HiSeq 2500, NextSeq 500, 1 × 50bp, 2 × 100bp, 2 × 75bp). Files from the sequencer were processed using a custom pipeline written in Snakemake^20^. The following preprocessing steps were performed. Reads from multiple lanes per sample were grouped together and basic QC performed using FastQC; low quality bases and adapter sequences were trimmed with Trimmomatic^21^; overlapping mate pairs merged into single reads with FLASH^22^; reads aligned to the UniVec^23^ core database with bowtie2^24^ to remove linkers, primers, adapters and vector sequences commonly used in the processing of DNA. The human component of these reads subtracted by aligning the cleaned reads to expanded reference human genome (GRCh38.p5, downloaded from ENSEMBL in January 2015) and removing all reads that mapped.

### Known infectome

The known infectome was characterized in a previously determined manner^9^. Briefly, non-human reads were aligned using BLAST^25^ against a database of known viruses, bacteria and fungi that had undergone masking for non-informative regions. These alignments were then used to estimate the abundances of taxonomic ids present in the database using GRAMMy^26^. These results were pushed to a database for later processing.

### Unknown infectome

Prior to assembly, an additional step of aligning reads to the NCBI nt database using BLAST resulted in the removal of an average of another 30% of reads as being “human-like” sequences. Following this, an iterative number of aggregated assemblies and cleanups were performed.

At any stage, the non-human reads of the sample(s) were assembled using SPADes^27^. Contigs that were identified as being plausible human sequences by alignment with BLAST (blastn, NCBI nt database), and those that were marked as being low-complexity using DUST^28^ were marked as ‘bad’ contigs. Following alignment (bowtie2) of the reads to the contigs, those reads that aligned to the bad contigs were removed prior to the next stage in the assembly process. This process began on a per sample basis, then went to the patient and cohort level. In addition, contigs from one level were aligned to the contigs of the next using LAST^29^ as a means of identifying the contigs deriving from the same genome and to perform quality control on the assembly process. The final set of contigs had coding regions predicted using PRODIGAL^30^, with the genes identified by homology to known or predicted protein coding sequences (blastx, NCBI nr database). Taxonomic position was determined by finding the majority consensus taxid based on the taxonomic lineage of each gene on the contig.Additionally, phage-like gene (taxids of 28883, 12333, 79205 present in the lineage) were ignored if this resulted in a more specific taxonomic assignment.

Novel contigs were selected by requiring a length >=1 kbp, coverage with BLAST below 20% of the contig length, gene coverage above 60% and average gene identity below 60%. The comparison with the HMP was achieved by downloading all assemblies from phases II and III of the project and aligning using LAST (lastal -q5 -e100) to a database built using the cohort contigs. Any alignment was taken as indicating homology. The Onecodex^14^ comparison was achieved by uploading a FASTA file of candidate novel contigs and downloading the kmer alignment table after aligning to their comprehensive Onecodex database. If coverage was >20%, the contig was excluded from later analysis.

To control for contamination from the column, a total of 275 million reads (2×75 bp) from six libraries made with either water or human DNA extracted from a cell-line were processed in the same manner as plasma samples. After sequencing, quality control and human removal performed in the same manner as the plasma samples above, they were aligned to the candidate novel contigs. The threshold to determine that a contig is a potential contaminant was set to an average one read fragment aligning per 500 bp of assembled contig. The nonhuman primate samples were also processed in a similar manner, and after quality control cleaning up and subtraction of likely primate sequences, they were aligned to the candidate novel contigs too.

All 125,842 sequences from a global metagenomic virome project^16^ were downloaded and aligned using LAST to the novel contigs. Those that had an alignment covering at least 50% of bases were selected. Metadata about the virome assemblies was obtained from supplementary table 19 of the publication, additional information obtained from JGI’s IMG/M ER.

As each cohort was assembled independently, there were contigs across cohorts that were likely to be derived from the same genome. Using LAST, the novel candidate sequences were aligned to themselves and those with more than 20% homology were grouped together and assembled using the default settings of CAP3^31^. Genes and their homologies were predicted on these clustered contigs in the same manner as above.

The solar system plot of contigs and their taxonomic assignment was constructed by first building a taxonomic tree using the ETE3^32^. The assignment of a contig to a given position on the tree was based on the majority taxonomic assignment of homologous genes present, ignoring phage-like genes if this made the assignment more specific. The tree initially included all levels of the lineage but was trimmed to major phylogenetic levels. The arcs over which a given contig was assigned was determined by the number of samples that were assigned to each phylum, with colors determined by both kingdom and phylum.

The anellovirus tree was constructed directly from the Newick tree produced by the MUSCLE^33^ (multiple sequence alignment program) web-service on the EBI’s website using the default settings, with sequences from the selected novel contigs and the reference anellovirus sequences downloaded from NCBI (accessions: NC_014078.1, NC_027059.1, NC_025966.1, NC_014092.2, NC_028753.1, NC_026663.1, NC_026662.1, NC_026138.1, NC_014070.1, NC_026765.1, NC_026764.1, NC_026664.1, NC_014480.2, NC_025727.1, NC_025726.1, NC_024908.1, NC_024891.1, NC_024890.1, NC_020498.1, NC_012126.1, NC_015783.1, NC_015212.1, NC_014097.1, NC_014096.1, NC_014095.1, NC_014094.1, NC_014093.1, NC_014091.1, NC_014090.1, NC_014089.1, NC_014088.1, NC_014087.1, NC_014086.1, NC_014085.1, NC_014084.1, NC_014083.1, NC_014082.1, NC_014081.1, NC_014080.1, NC_014079.1, NC_014077.1, NC_014071.1, NC_014069.1, NC_014076.1, NC_014075.1, NC_014074.1, NC_014068.1, NC_014073.1, NC_014072.1, NC_009225.1, NC_007014.1, NC_007013.1, NC_002076.2, NC_002195.1).

Codon based classification was performed by first computing the codon frequency for all genes from the novel, divergent and known contigs. Three AdaBoost classifiers^34^ built using 128 decision trees were trained on the genes with identified superkingdoms that had >80% identity, 20-80% identity, and >20% identity. The remaining genes then had their superkingdom predicted based on these trained models, with the majority consensus determining the assigned superkingdom. Then all genes from a given unrooted contig were grouped and the superkingdoms of the gene assignments used to determine the superkingdom of the contig.

### Validations

The other cfDNA samples were downloaded from the Sequence Read Archive (SRA) using sra-tools and processed through the pipeline in the same manner as all the other samples. The reads remaining after the second round of human subtraction were aligned using bowtie2 to the database of potential novel contigs. Accessions for the single-stranded cfDNA preparation of patient L79 are: SRR3067214, SRR3067215, SRR3067216, SRR3067217, SRR3067218, SRR3067219, SRR3067220 and SRR3067221. The accessions for the other maternal cfDNA samples are: SRR1705787, SRR1705788, SRR1705789, SRR1705790, SRR1705791, SRR1705792, SRR1705793, SRR1705794, SRR1705795, SRR1705796, SRR1705797, SRR1705798, SRR1705799, SRR1705800, SRR1705801, SRR1705802, SRR1705803, SRR1705804, SRR1705805, SRR1705806, SRR1705807, SRR1705808, SRR1705809, SRR1705810, SRR1705811, SRR1705812, SRR1705813, SRR1705814, SRR1705815 and SRR1705819.

Primers were designed using the primer3^35^ bindings for python [https://libnano.github.io/primer3-py/]. Default settings were used except for: the primer product size range for between 60-100 nt; the minimum primer temperature of 59C; a maximum primer temperature of 61C; and the number of primers to return for each sequence of 20. After primers were designed for 700 novel contigs seen most often in samples, each primer sequence (typically 20 nt) was aligned to the NCBI nt database using BLAST (method=”blastn-short”, evalue=1000) to filter out primers that may target known sequences. Up to four mismatches were tolerated (equivalent to 80% alignment coverage) for each end of the primer and those that aligned to the same sequence were filtered out. An additional filter to remove primers that targeted similar regions was also employed. For the contigs of interest primers were ordered from IDT. Libraries containing suspected sequences were thawed and 1 uL was diluted in 15 uL of water, followed by additional dilution if this minimized artifacts from too-high loadings. Amplification was performed using the standard protocol for NEB Phusion PCR Master Mix, with 20 uL in each reaction chamber used. The reference PCR cycle was at performed at 60C for 30-35 cycles. E-Gel EX 2% agarose kits (ThermoFisher) were used to test for amplification of the template. A 10bp or 25bp ladder (ThermoFisher) was used to quantify the sizes of the template and if they were of the correct size as designed for.

## Acknowledgments

This work was supported by…was supported by…We would like to thank David Grimm and Helen Luikart for providing us with extra plasma samples for validation experiments.

## Author contributions

R.W, S.K., Y.E., Y.B., D.K.S and G.S. were involved in patient recruitment and sample collection. J.C., I.D.V, W.K., W.P., L.M., N.N and J.O. were involved in extracting nucleic acids from samples and preparing sequencing libraries. M.K., Mi.K., I.D.V. and S.R.Q conceived of the the research idea. M.K performed analysis. M.K., N.D.W and S.R.Q wrote the manuscript.

The authors declare no competing financial interests.

Correspondence and requests for materials should be addressed to

---

**Figure S1.**
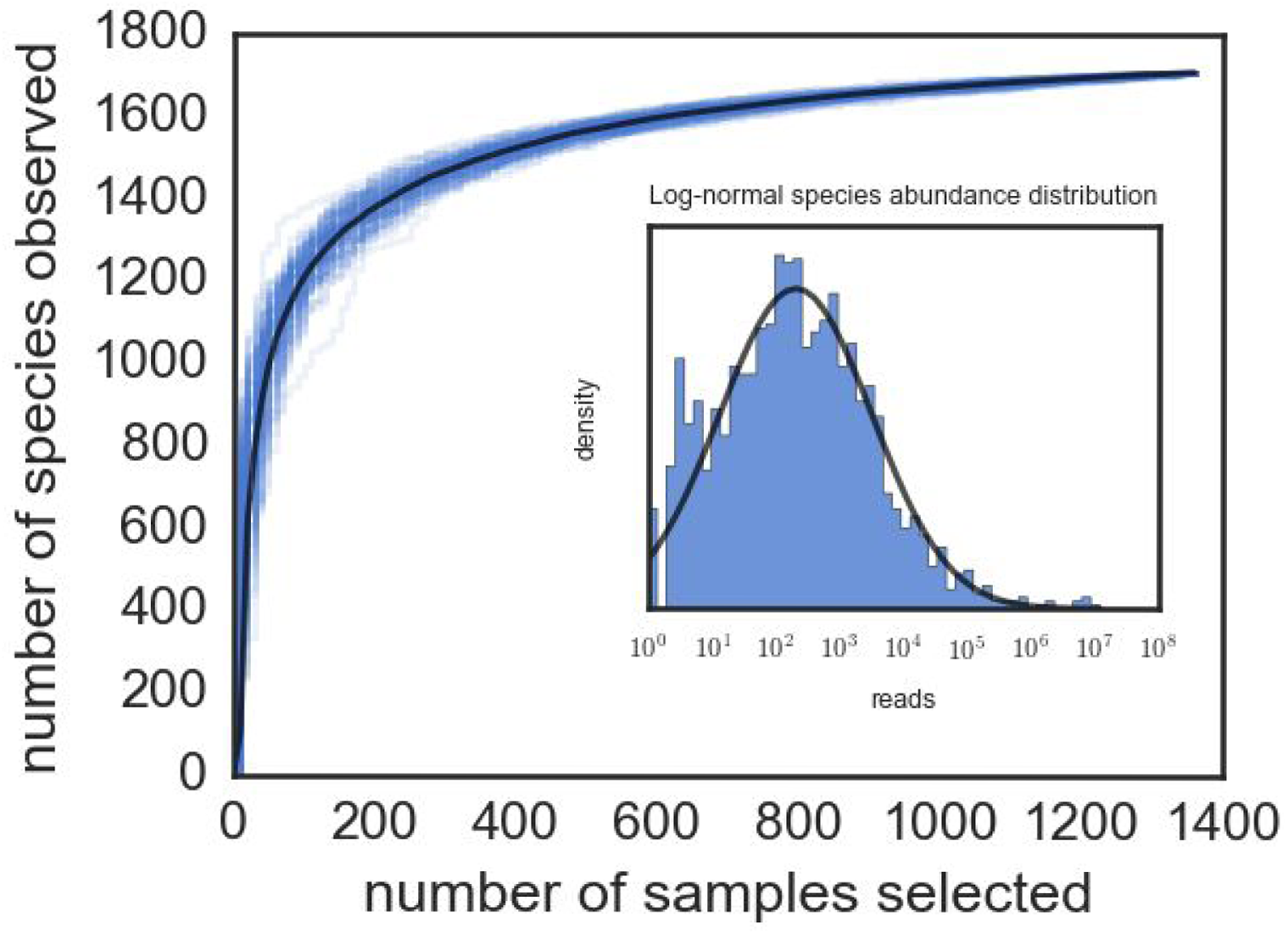
Rarefaction curve of the number of species observed in downsampled sets. Black line is the mean of 100 different subsamples. Inset: species abundance distribution showing the number of reads aligning to each species, the log-normal distribution fitted using the sample mean and standard deviation.

**Figure S2.**
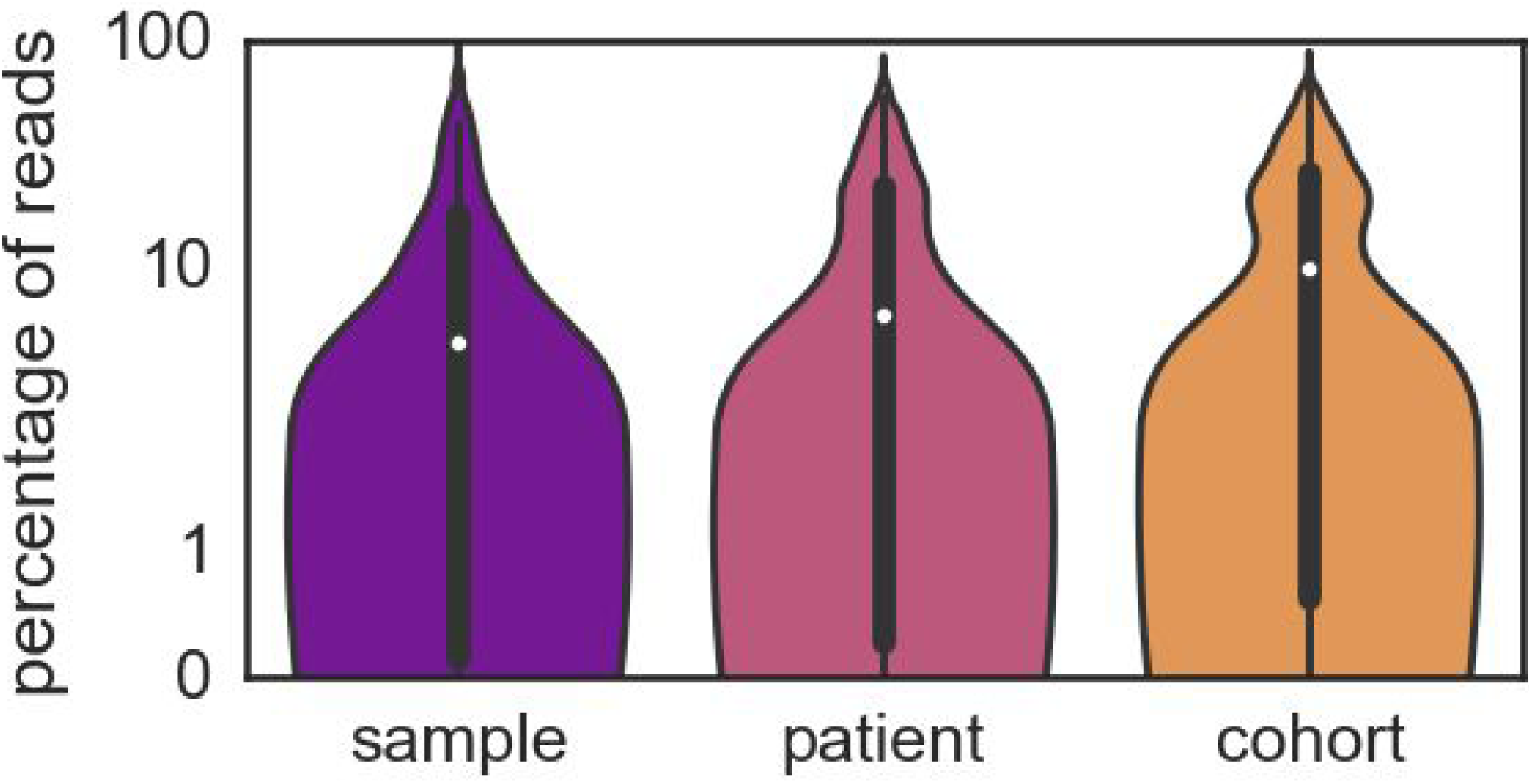
Violin plots of percentage of reads aligning to each stage of assembly. There is an upward trend of the median and interquartile range as more samples were pooled in the assemblies.

**Figure S3.**
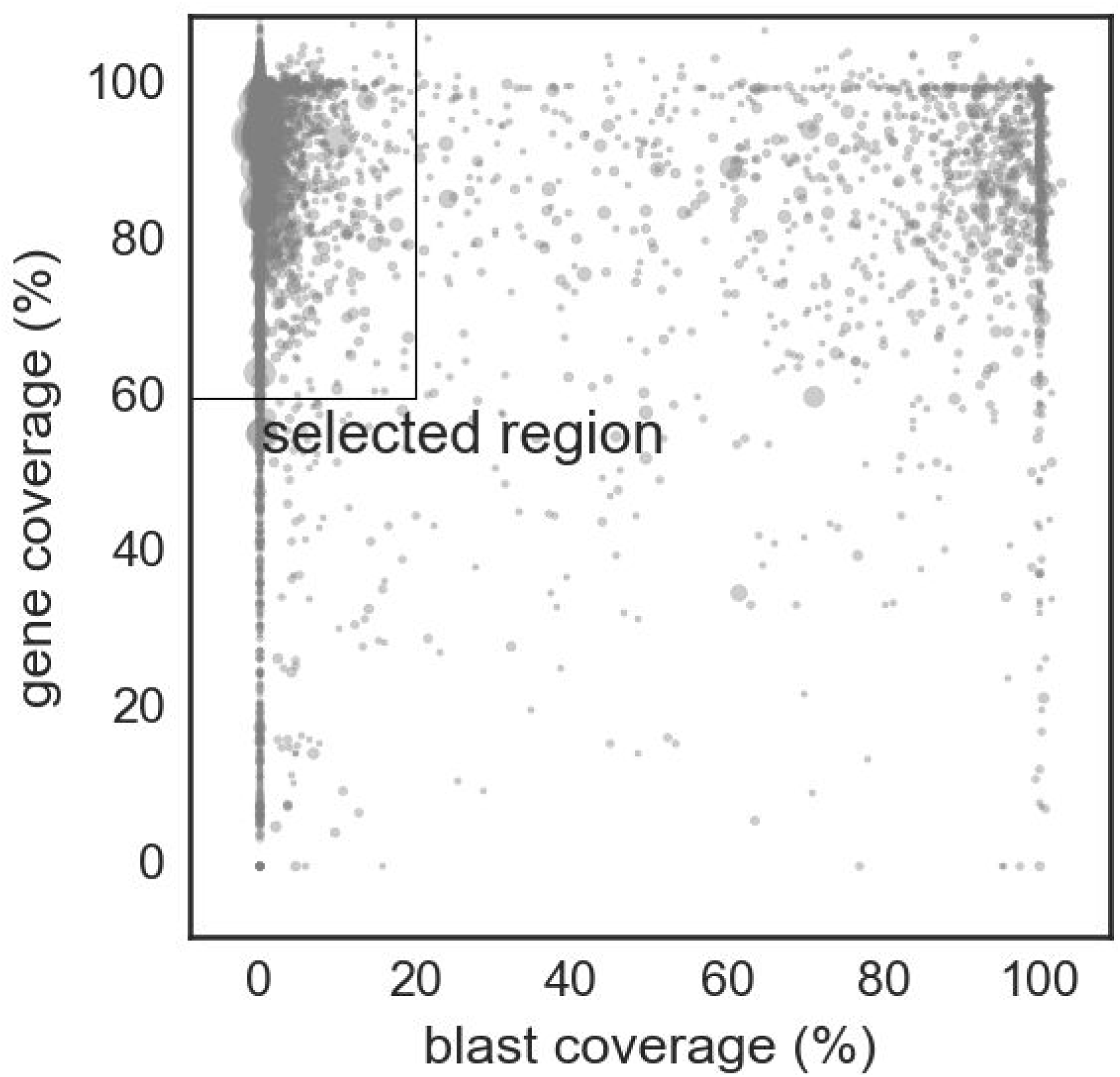
All contigs at the cohort level (>1 kbp) showing one of the filters for selecting for novel candidates: the blast coverage must be below 20% and the gene coverage above 60%. Size of circles is proportional to contig length.

**Figure S4.**
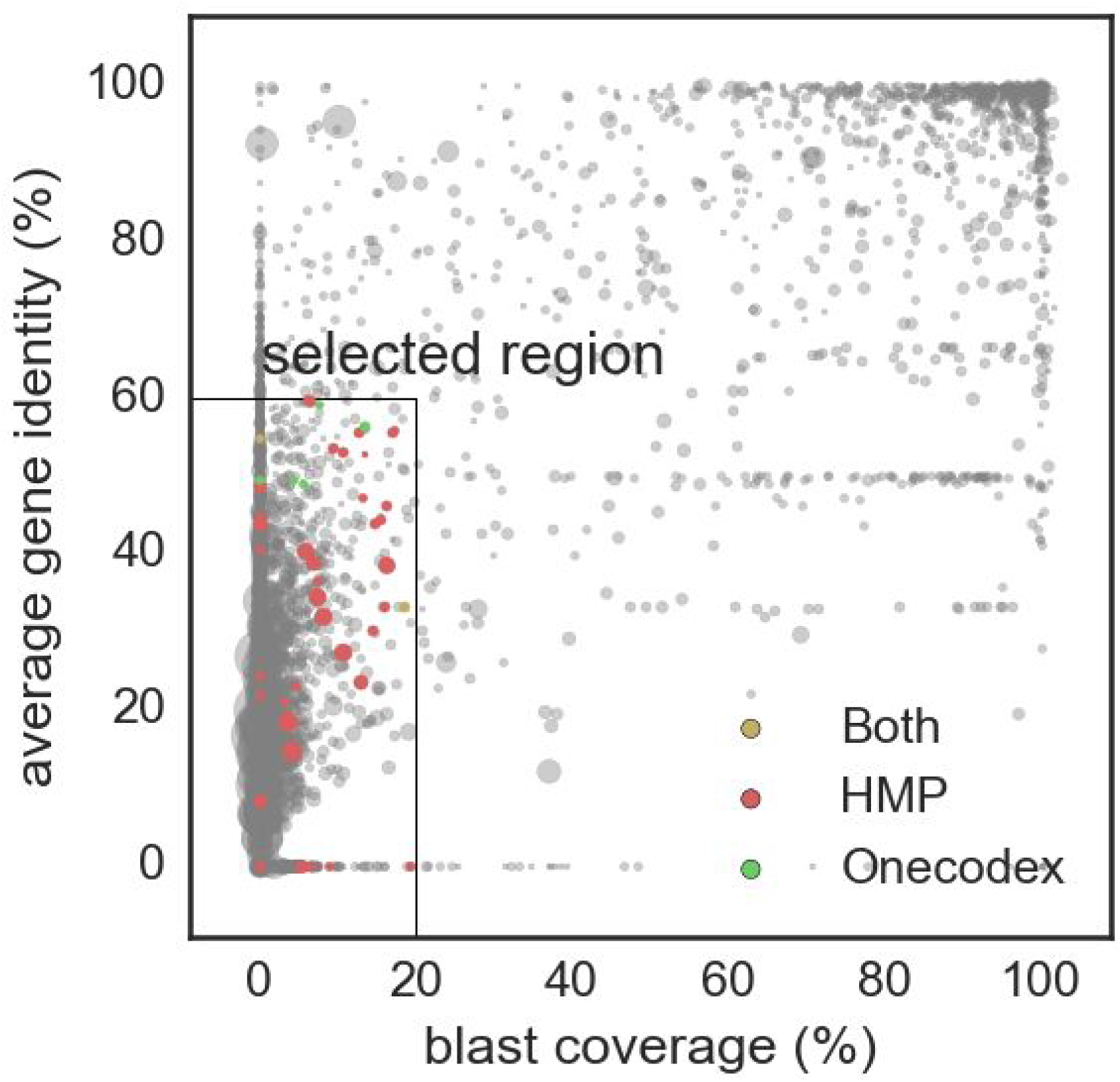
All contigs at the cohort level (>1 kbp) showing the second main filter for selecting for novel candidates: the additional requirement that the average gene identity on a contig is <60%. Size of circles is proportional to contig length. Coloured circles are additional contigs filtered due to having homology in HMP dataset, Onecodex’s database, or both.

**Figure S5.**
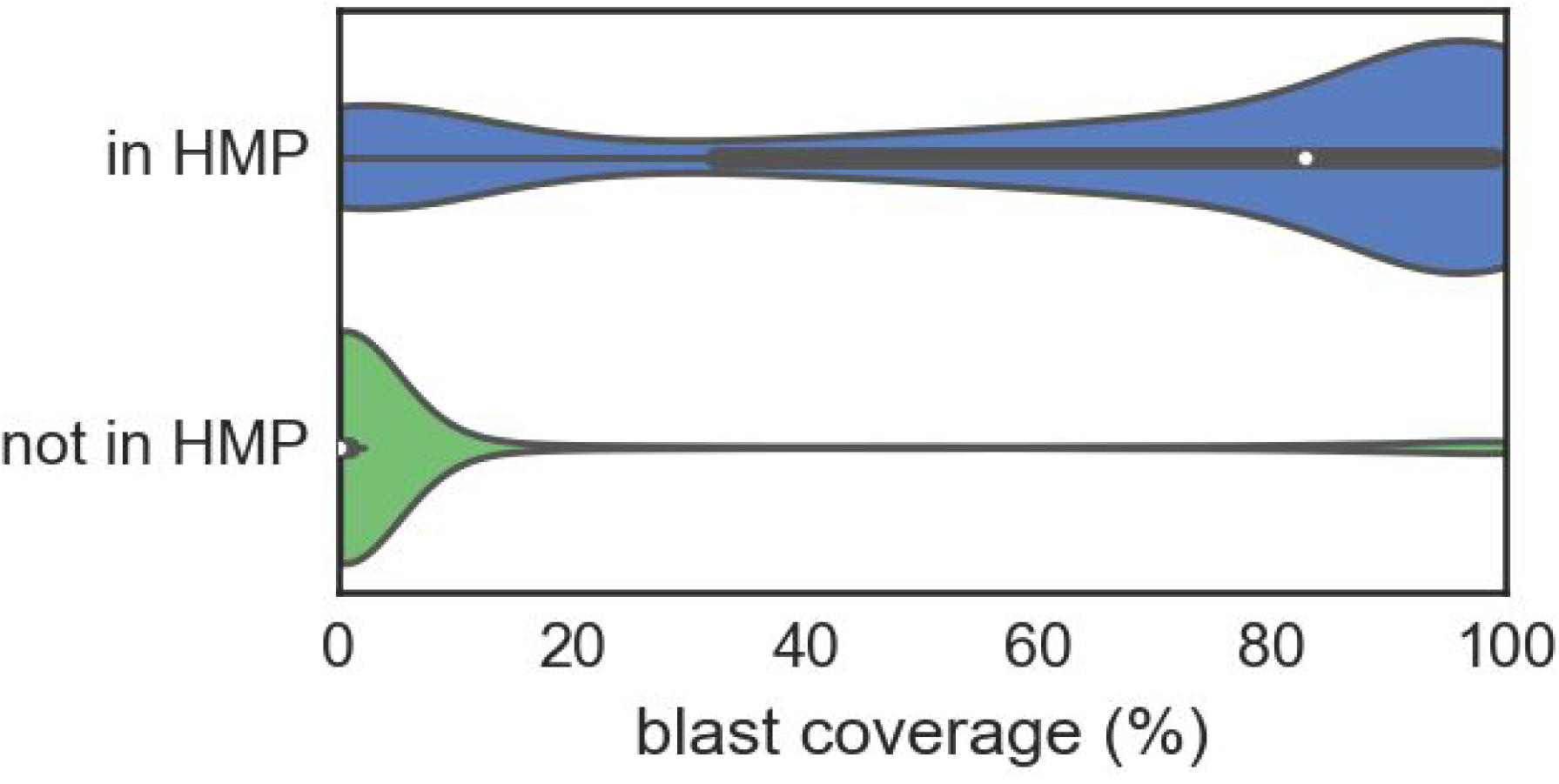
Violin plots showing the distribution of blast coverage of contigs partitioned by whether they are observed in the HMP data. The majority of those in HMP have blast coverages above 80%, whereas those not seen in HMP have <10%.

**Figure S6.**
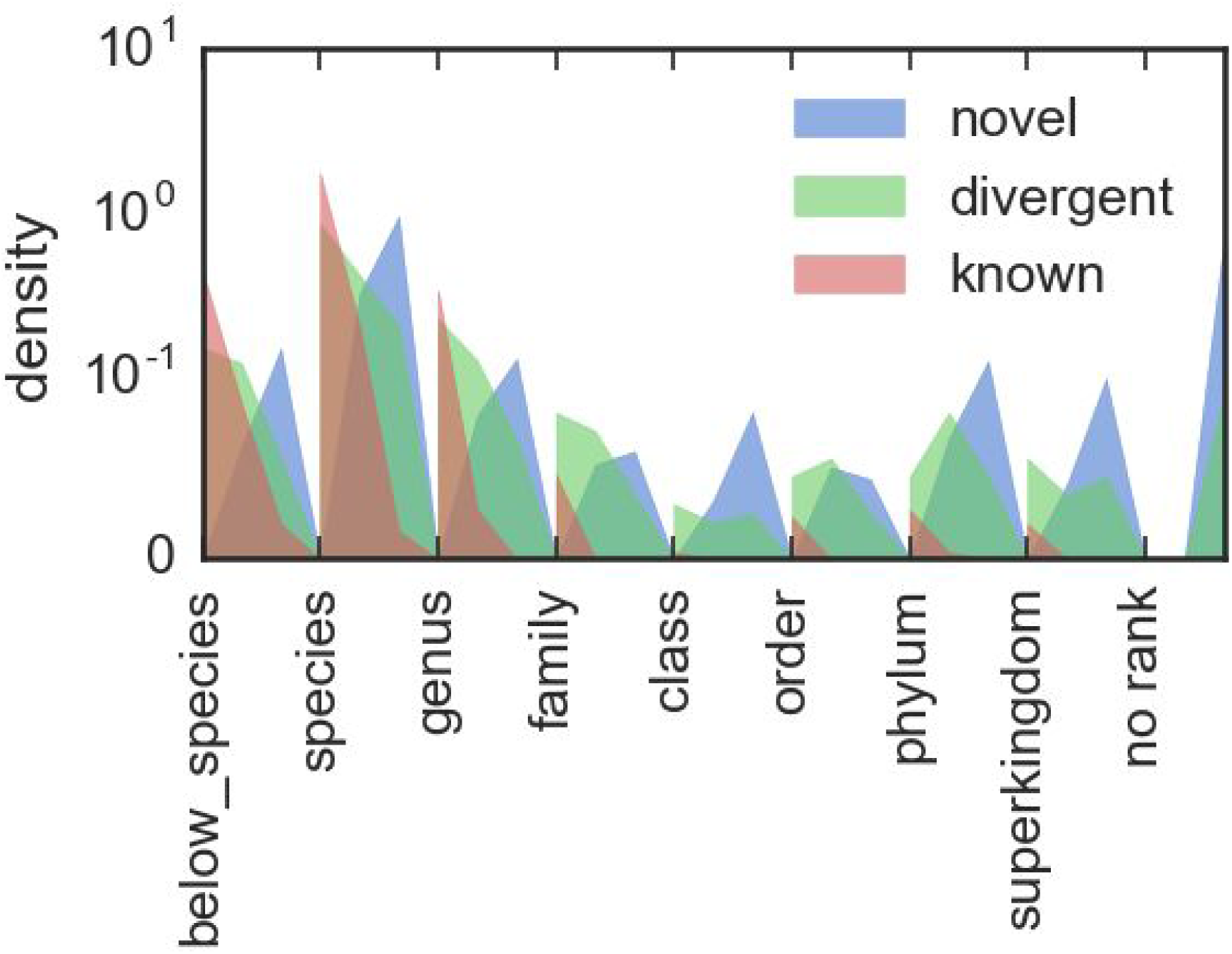
Are plots of the assigned taxonomic ranks of novel, divergent and known contigs as presented in the solar system plots. Each rank is broken up into three sublevels, corresponding to average gene identities of 100-67%, 66-34% and 33-0%. This allows both the overall differences in rank assignments to be compared between the contig types, and the difference of average gene identity within each rank to be contrasted.

**Figure S7.**
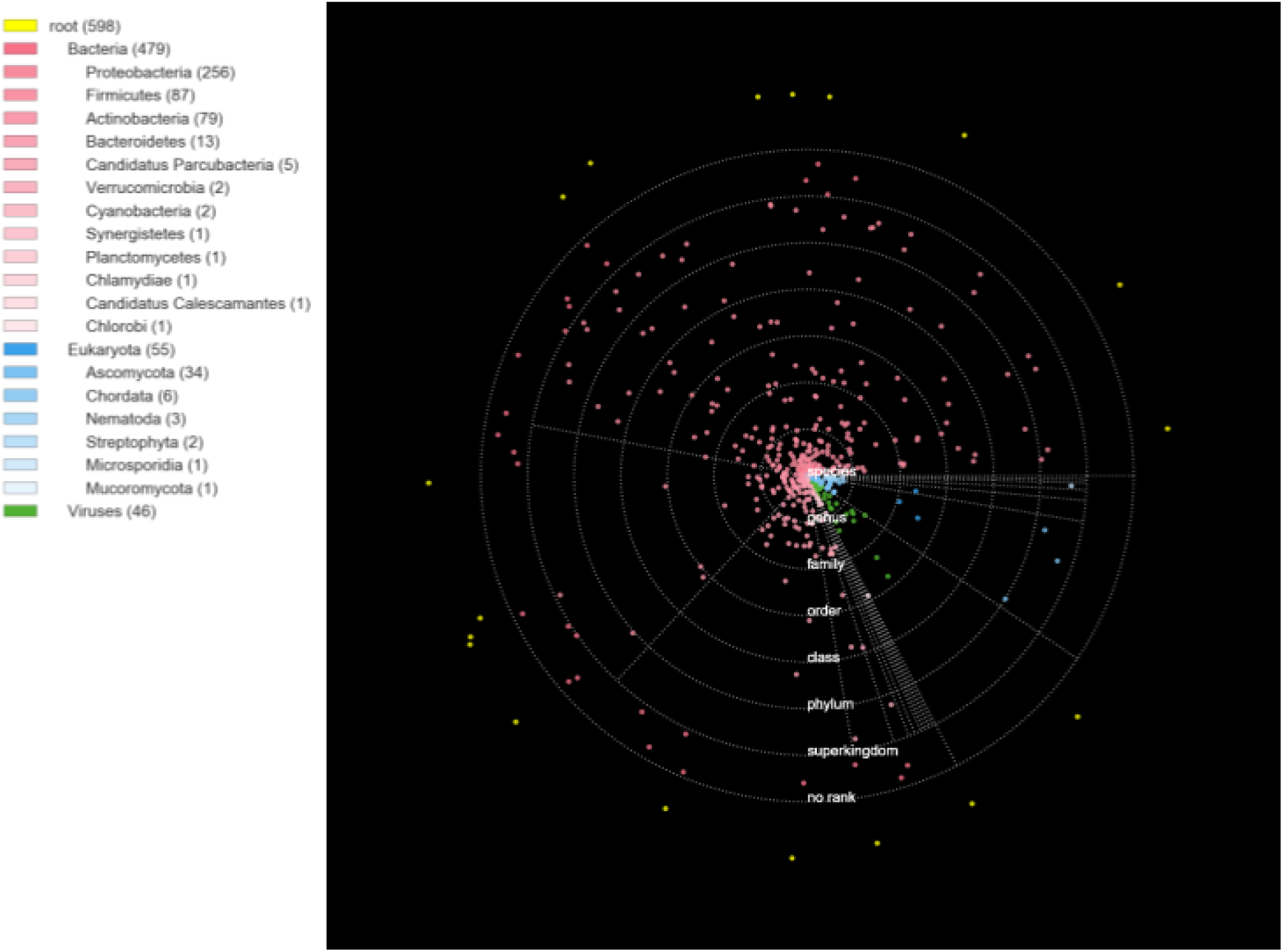

Solar system plot for divergent contigs

**Figure S8.**
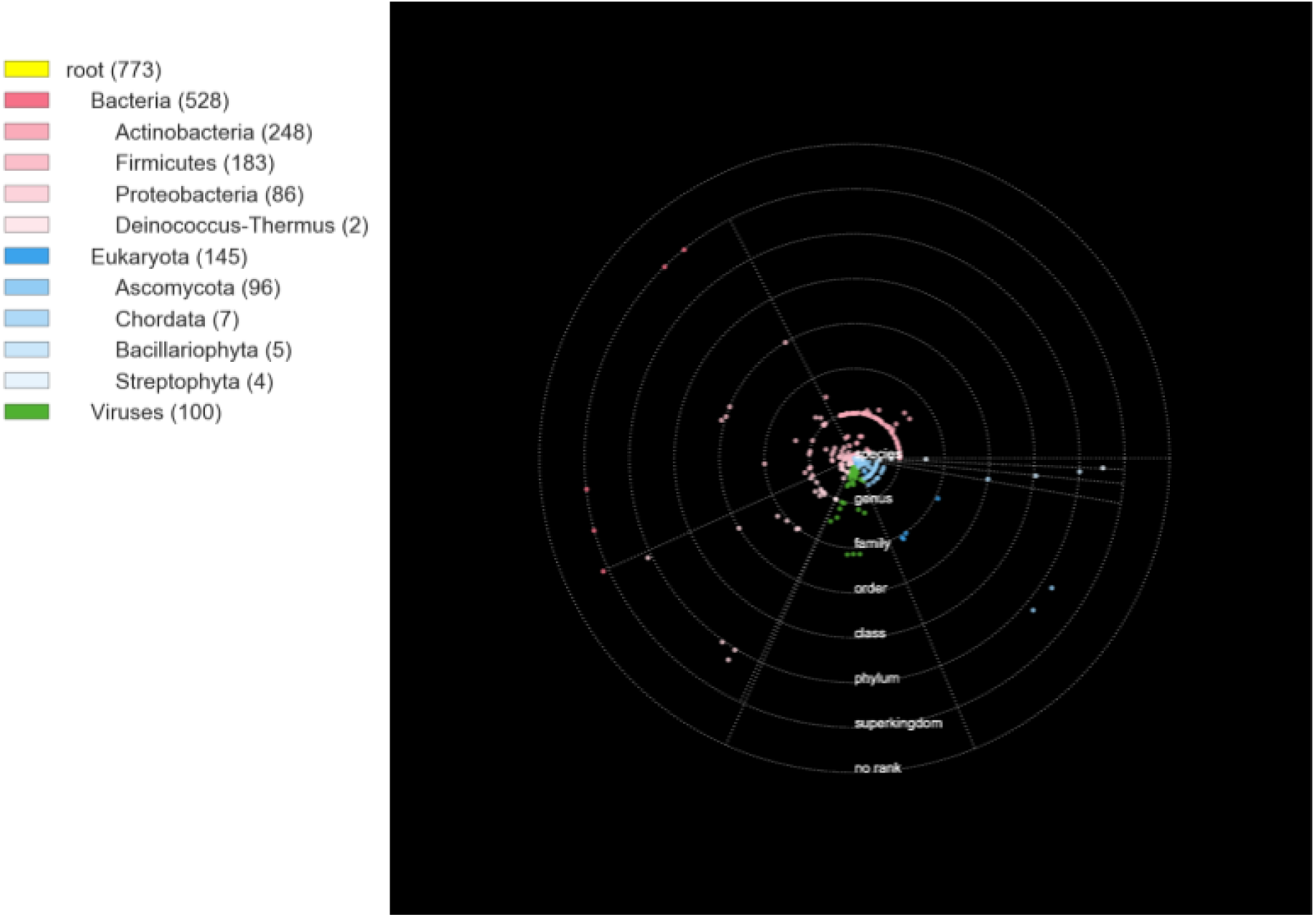

Solar system plot for known contigs

**Figure S9.**
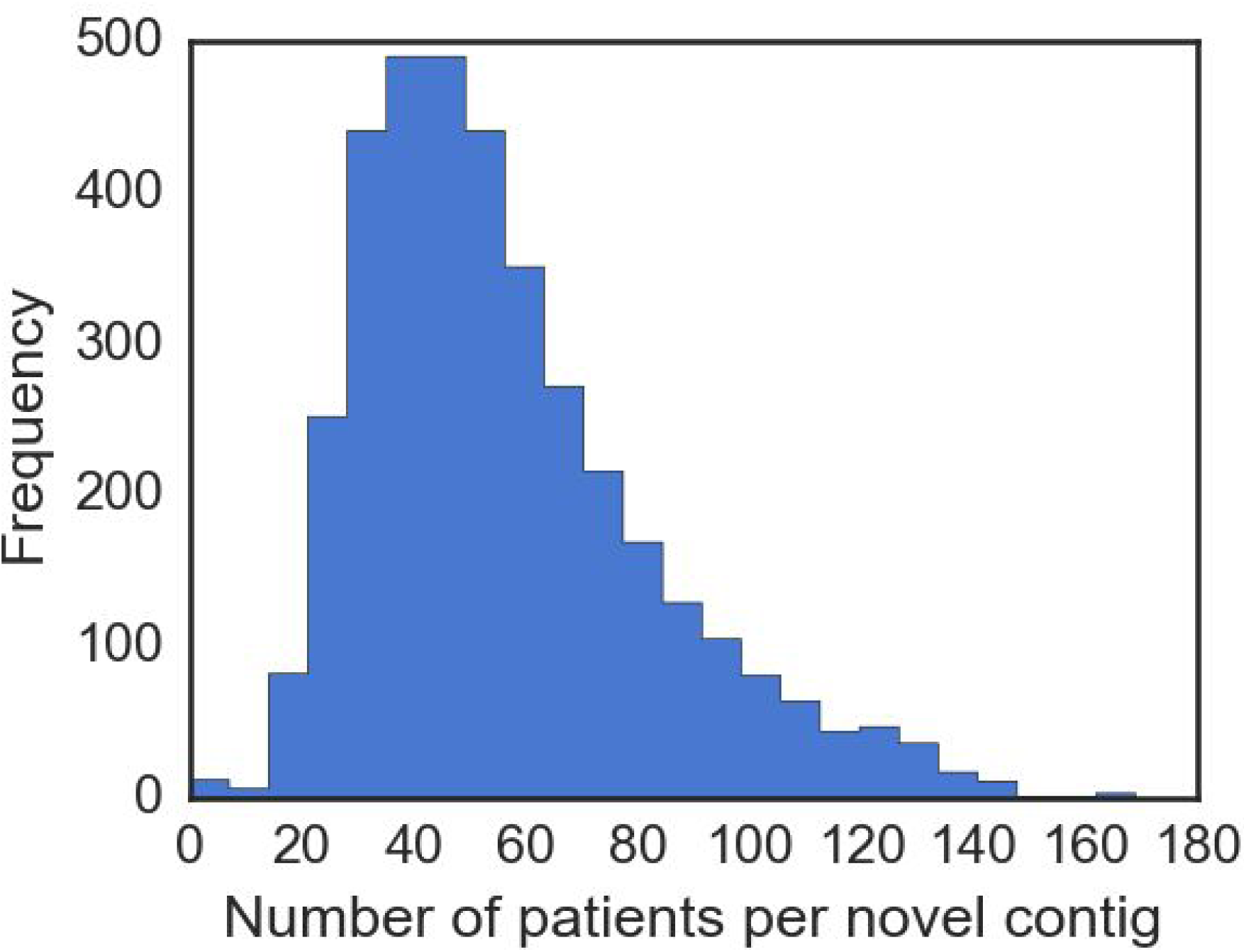
Histogram showing the number of patients (total of 188 in this study) each novel contig was observed in.

**Figure S10.**
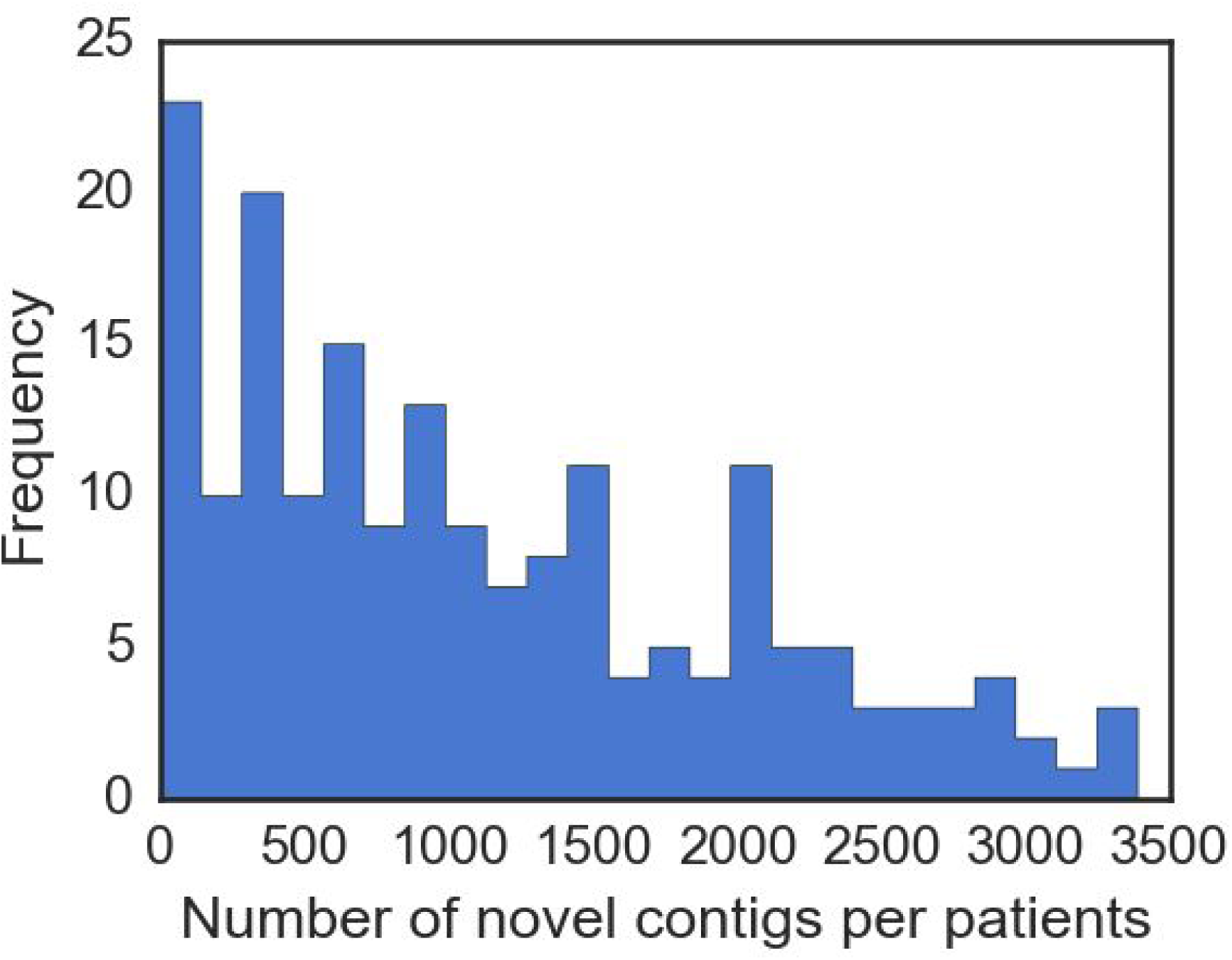
Histogram showing the number of novel contigs (total of 3761) observed in each patient.

**Figure S11.**
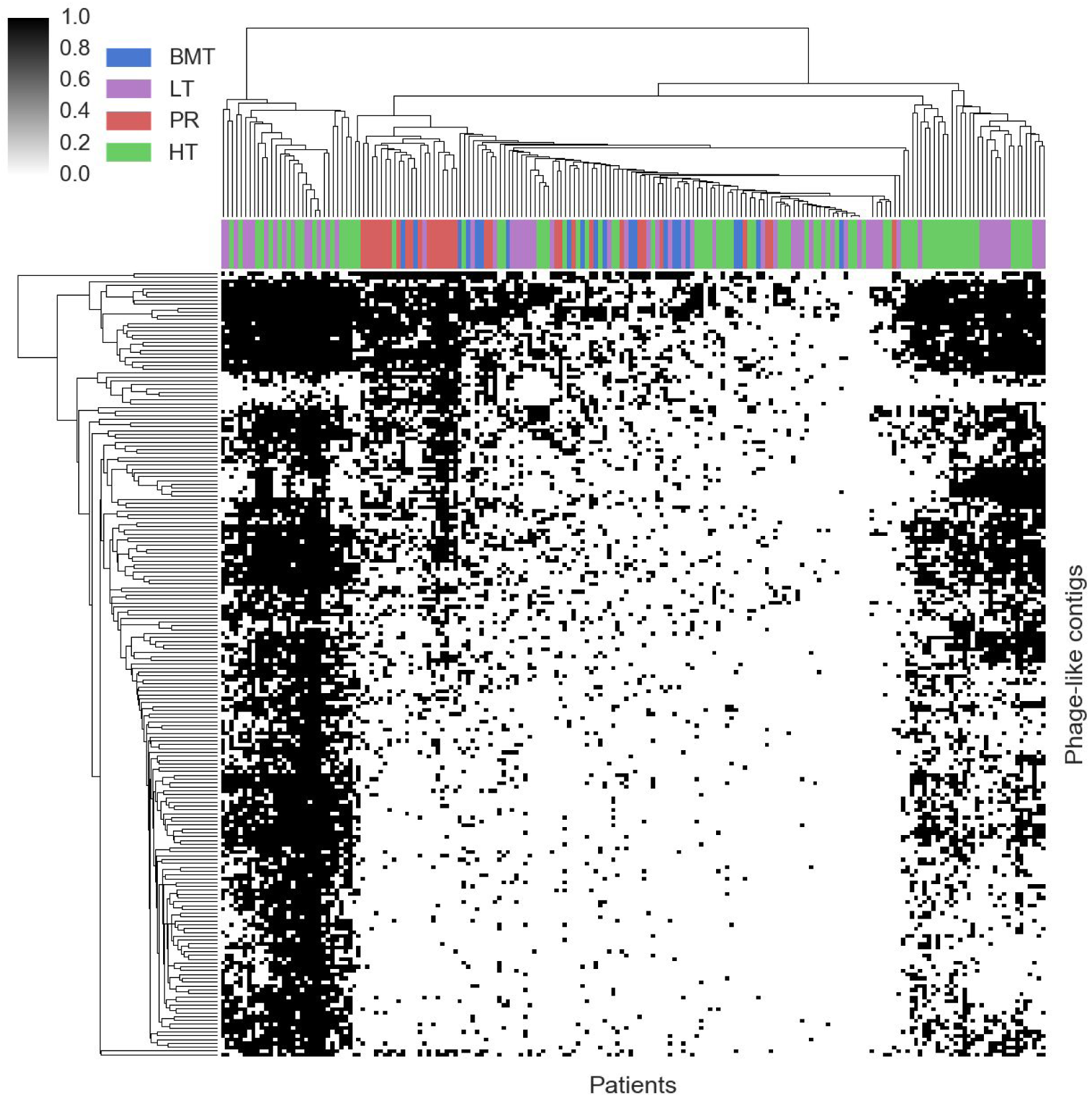
Heatmap showing the presence (black) or absence (white) of phage-like contigs across all patients. There is no strong cohort specific clustering.

**Figure S12.**
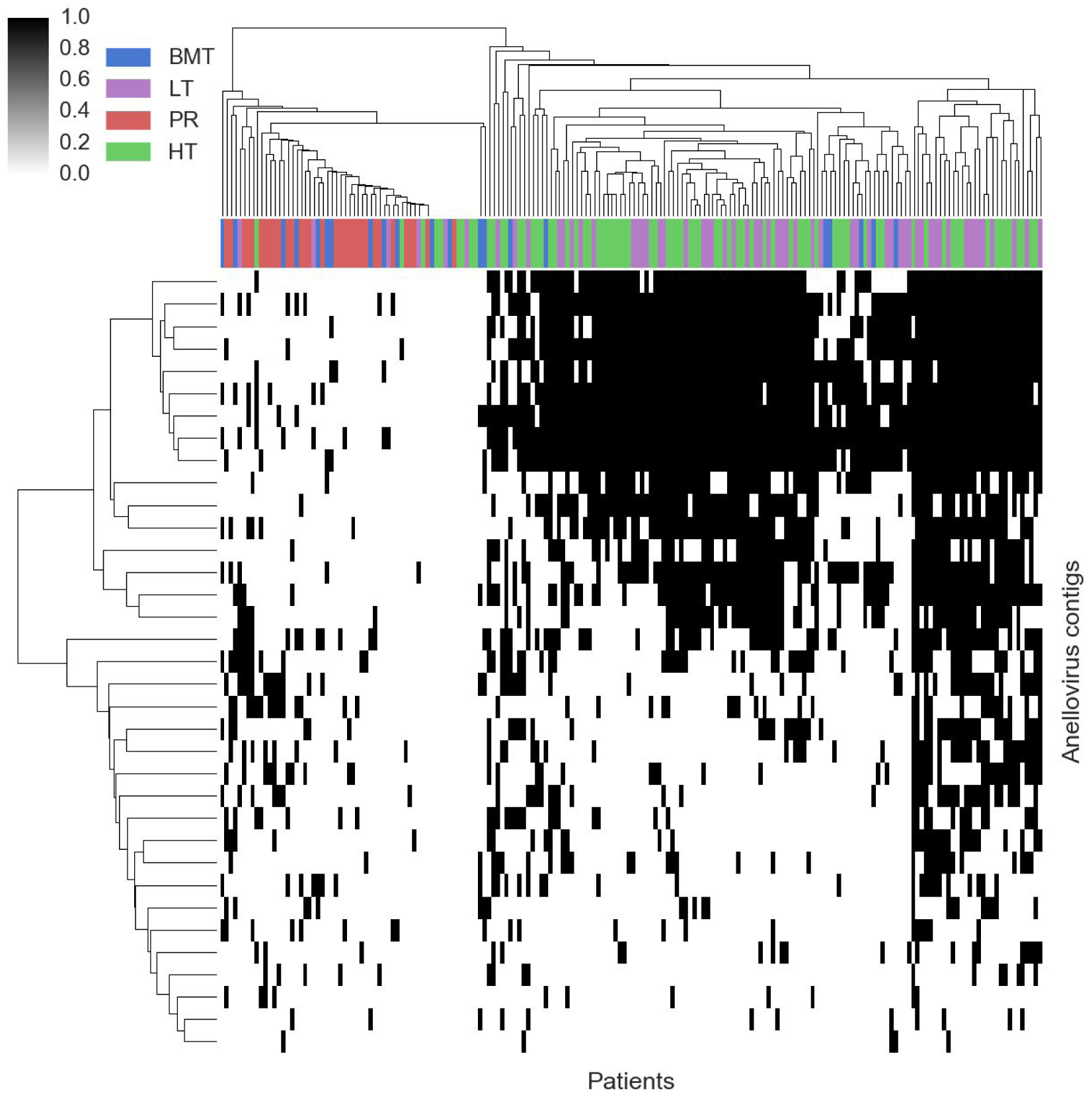
Heatmap showing the presence (black) or absence (white) of anellovirus contigs across all patients. There is strong cohort specific clustering between immunocompromised (heart and lung transplant) and others (bone marrow transplants and pregnant women).

**Figure S13.**
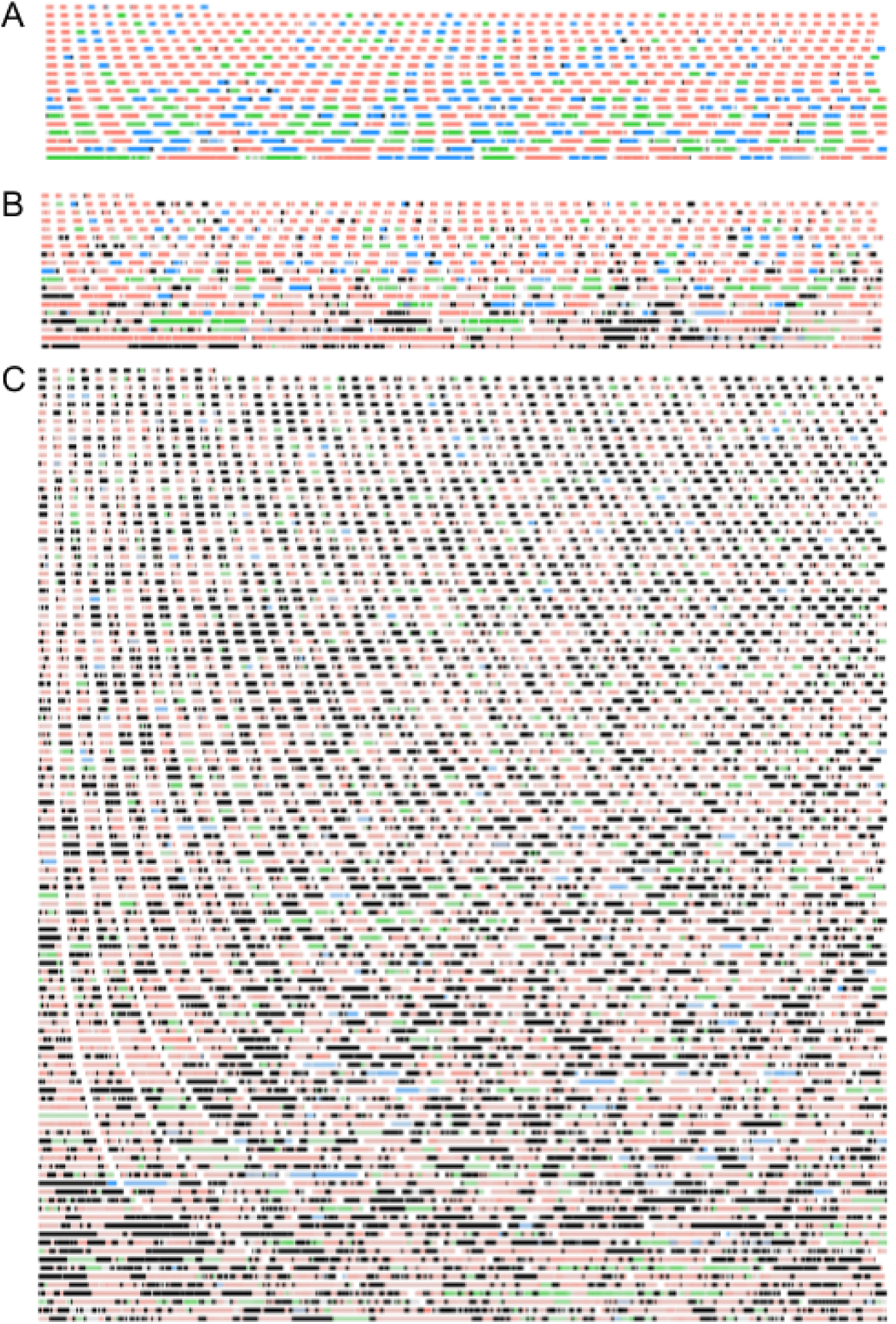
Plot of the all assembled contigs (>1 kbp) in the categories of A) known B) divergent and C) novel.Colours indicating gene superkingdom (salmon: bacteria, viruses: lime green, eukaryota: dodger blue, archaea: fuchsia, no homology: black, no gene: grey.

**Figure S14.**
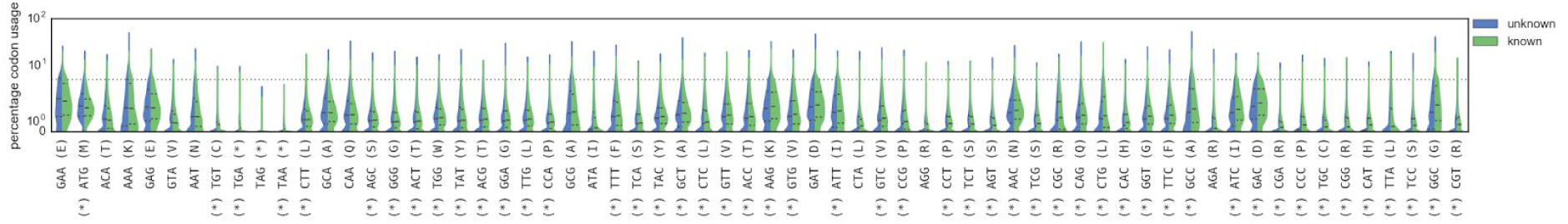
Violin plots of codon usage, stratified by whether the gene had homology (known) or not (unknown). Codons are ordered from left to right by the difference in the median usage in each of the two classes. Starred codons indicate a significant difference in distributions (p < 1e-3 after Bonferroni corrections, Mann-Whitney U test).

**Figure S15.**
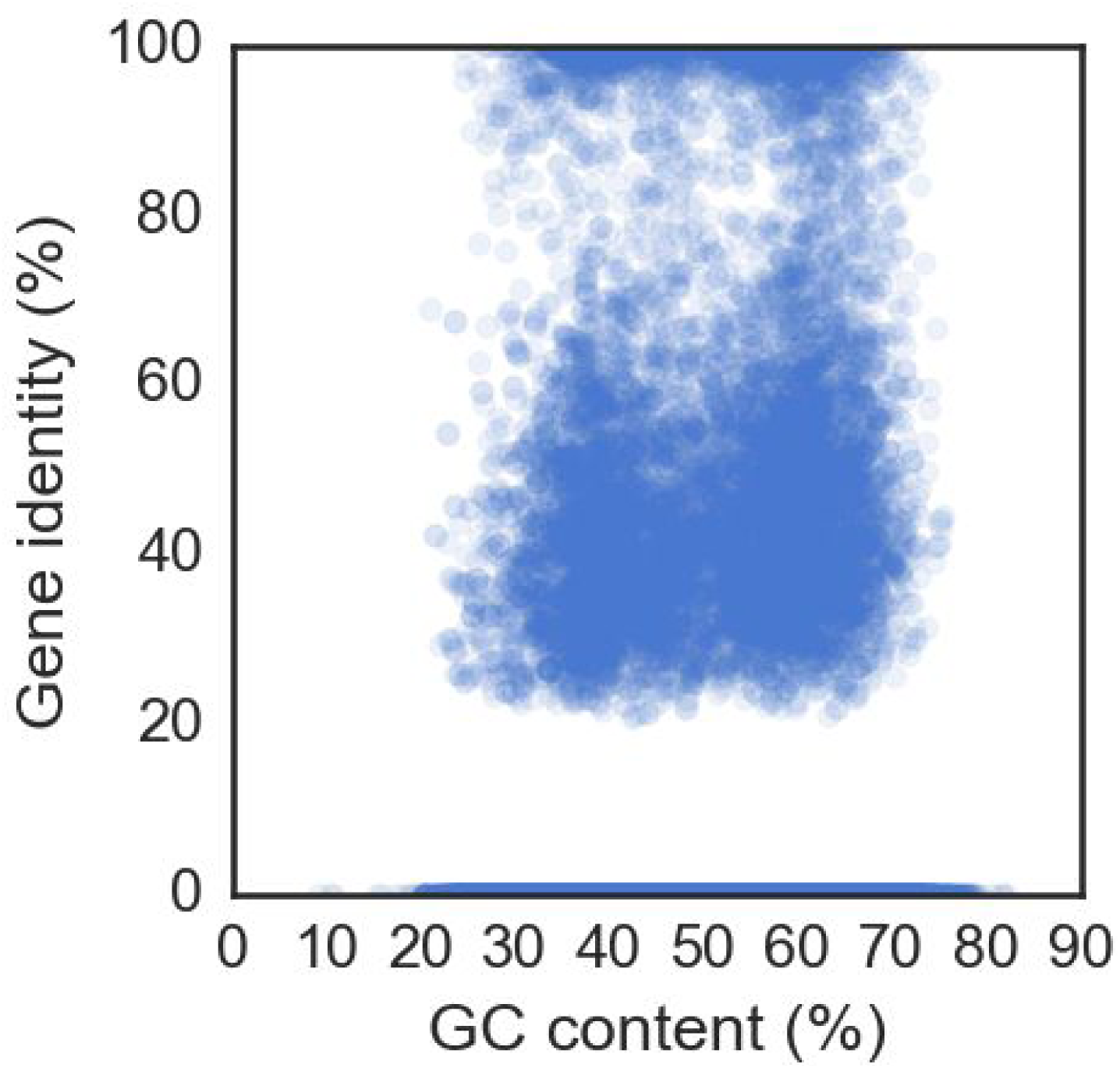
Scatter plot of the of all genes comparing the GC content to the (if existent, otherwise 0%) homologous gene identity, showing no clear bias between them.

**Figure S16.**
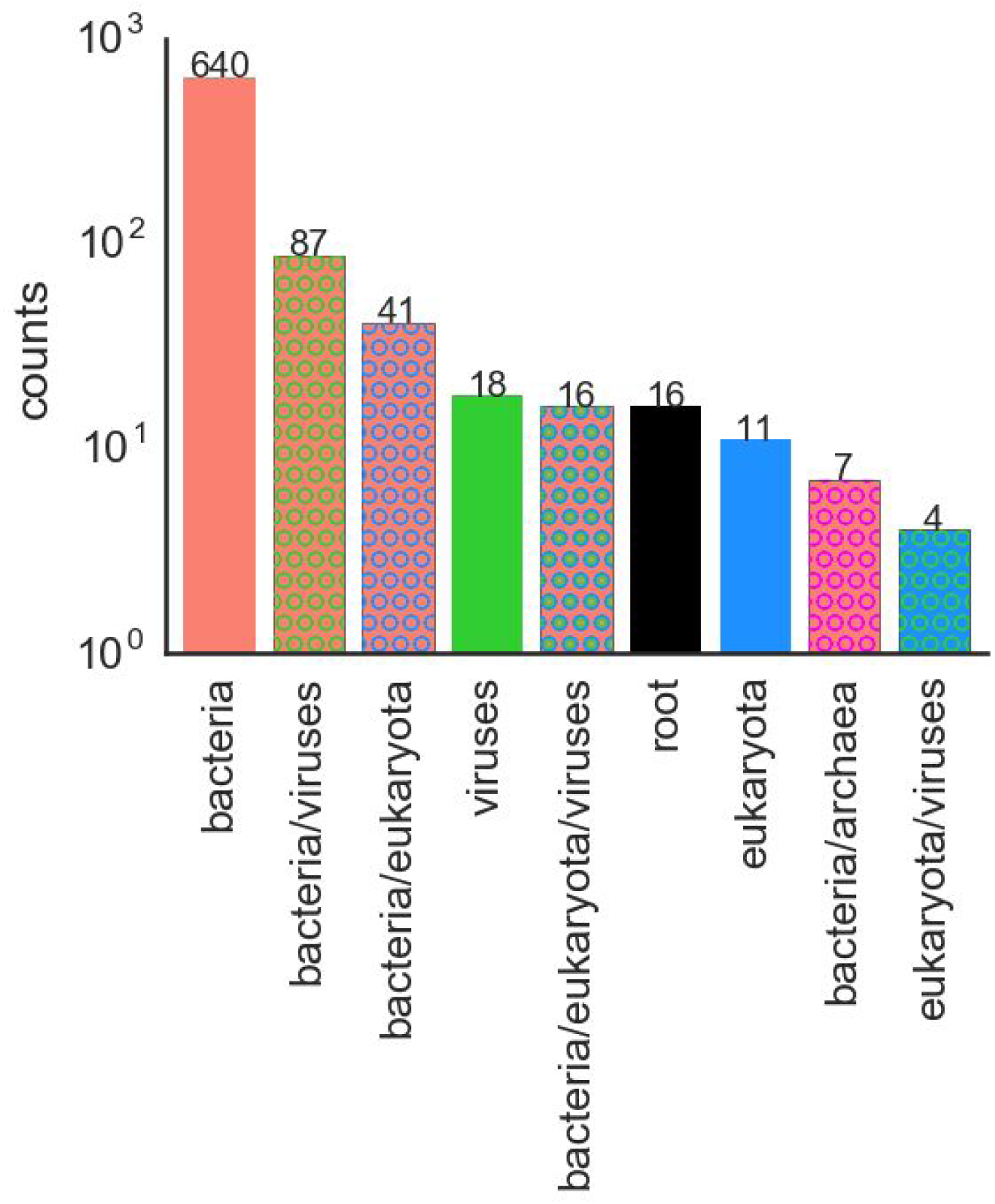
Potential assignments of the 840 novel contigs that have genes without any alignment-based homologies, based on similarities of codon usage to identifiable genes. Most are bacterial or phage-like, with a sizeable portion possessing genes that are associated with multiple domains of life.

**Figure S17.**
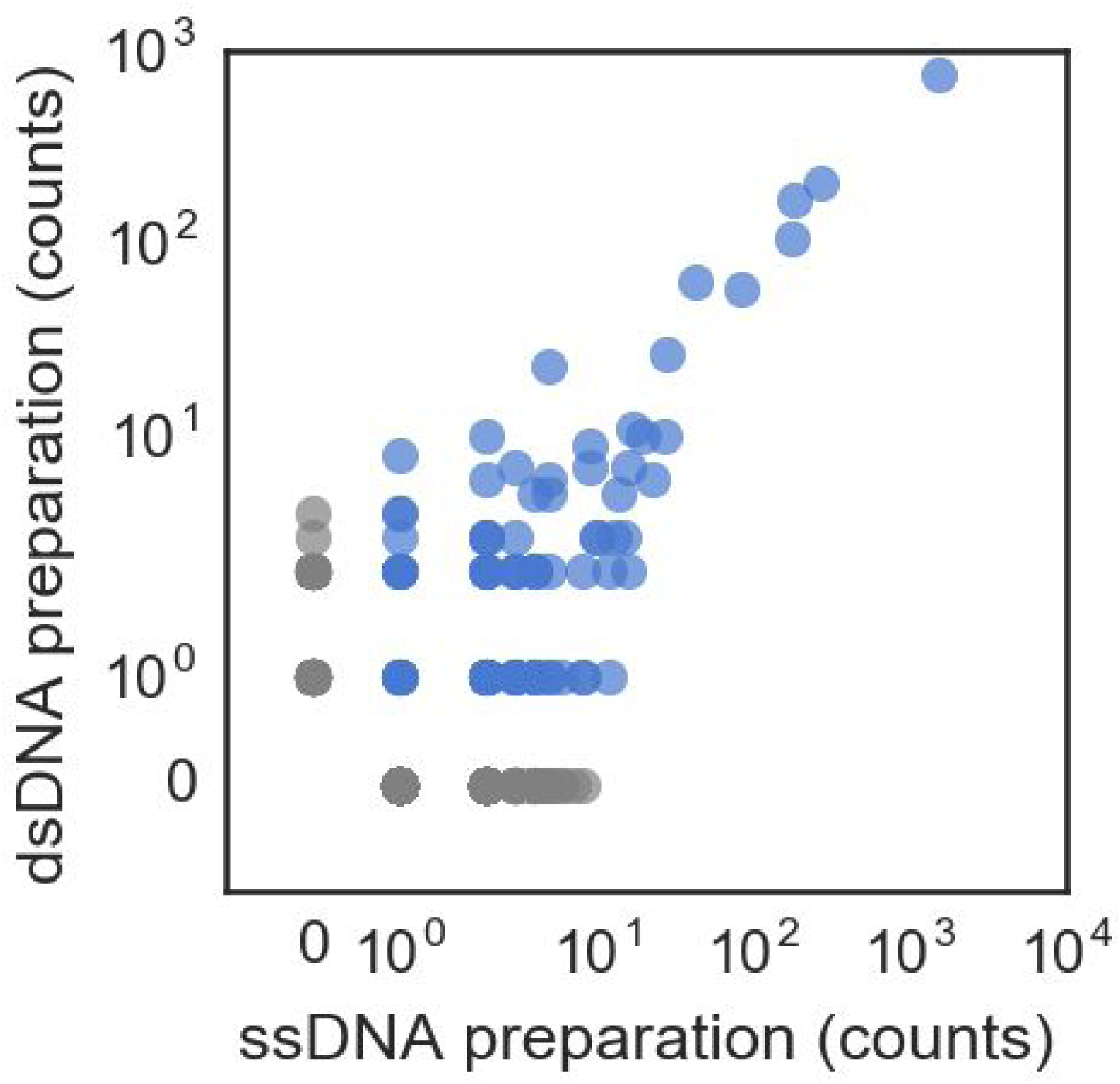
Scatterplot of the number of reads aligning to novel contigs for eight samples from patient L79. Besides a difference in absolute number, the relative rank of the most abundant novel contigs is concordant between our method (dsDNA preparation) and an alternative method (ssDNA preparation). Blue dots are contigs with counts in both preparations, grey dots are only observed in one.

**Figure S18.**
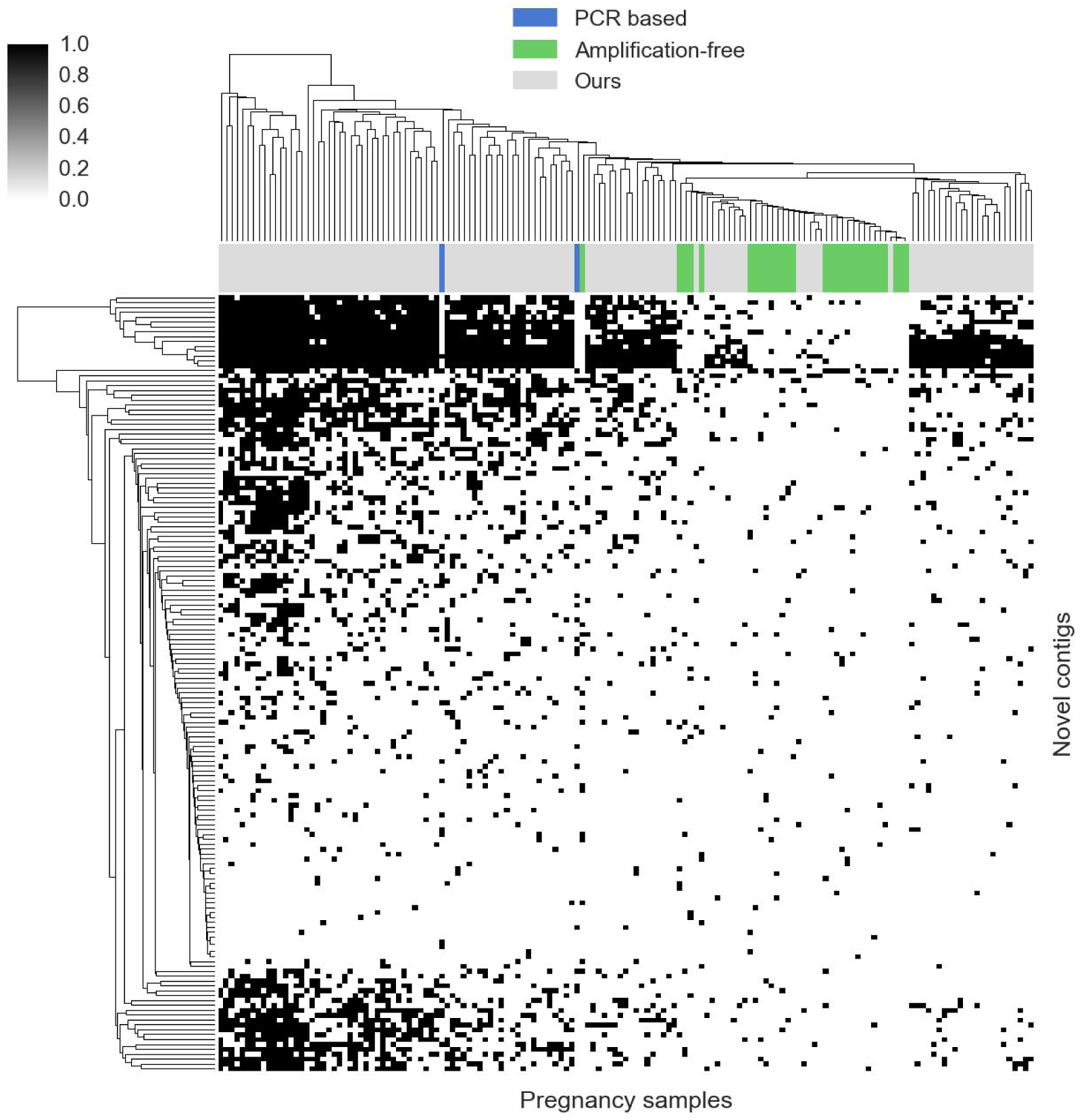
Heatmap of the presence of novel contigs in cell-free DNA samples from pregnant women. Coloured columns are from an independent lab and used a different protocol for extraction.

**Figure S19.**
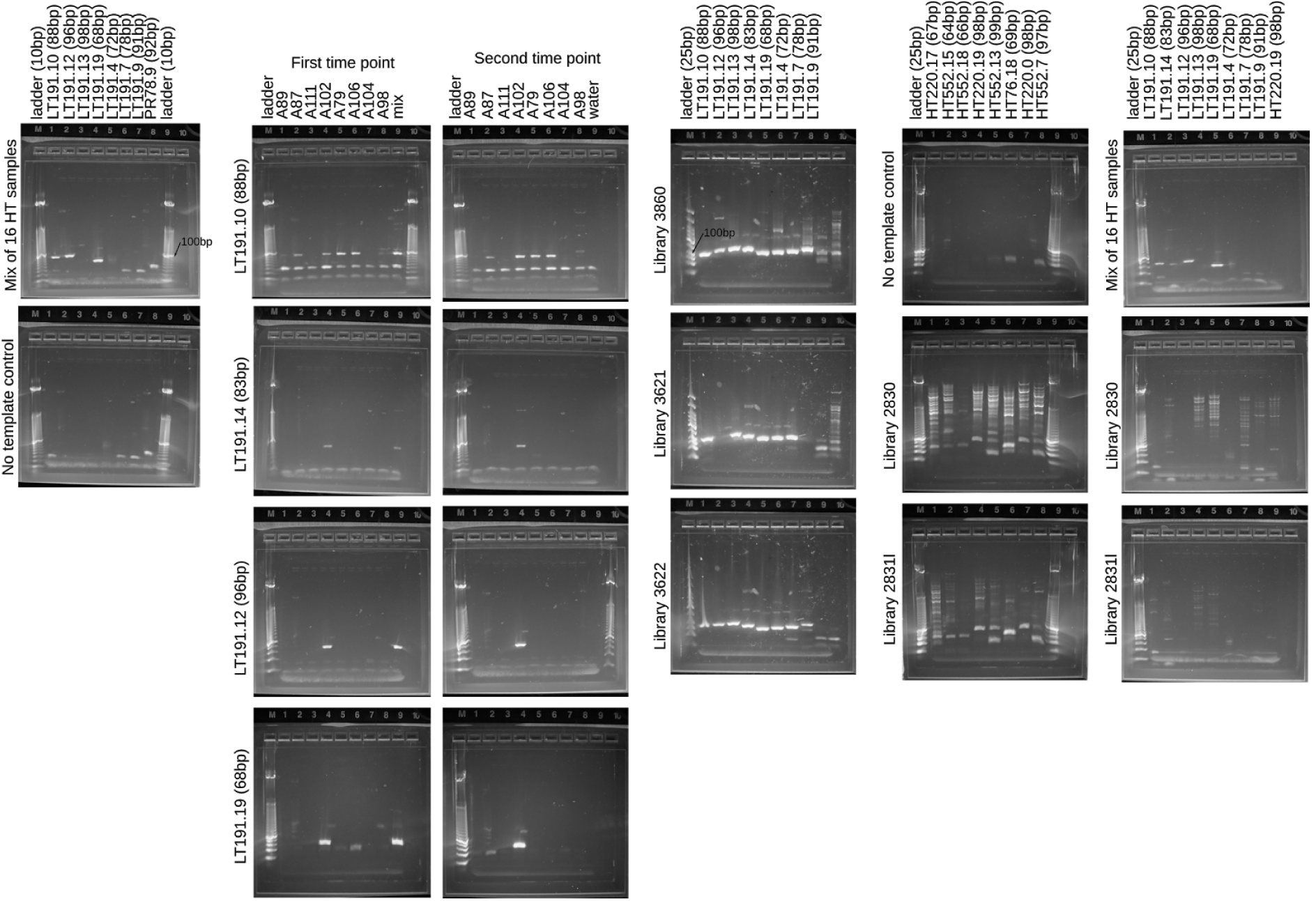
Left column: gels run to test for the presence of fragments of a novel torque teno virus in cfDNA extracted from 16 samples deriving from heart transplant (HT) recipients. Lower gel is the no template control. Right column: A slightly different panel of primers: upper is the same mix of 16 HT samples, the lower two panels used DNA derived from sequencing libraries from a heart (IL2830) and lung (IL2831) transplant recipients and have only a weak signal for LT191.14. The second and third column show the 16 samples individually for each of the four positively observed LT191 fragments. First and second time points typically differ by two-three months. The fourth column is the positive control for the novel TTV, the three libraries chosen were all known (from sequencing) to have high numbers of molecules aligning to these regions. The fifth column includes the templates from unrooted contigs, no size appropriate signal was seen.

